# Autozygosity and genetic load as sensitive early warnings of butterfly population decline

**DOI:** 10.64898/2026.08.05.742957

**Authors:** Sebastiano Fava, Marco Gargano, Dona Kireta, Paolo Gratton, Donatella Cesaroni, Alessio Iannucci, Claudio Ciofi, Roberto Biello, Marco Gerdol, Giorgio Bertorelle, Emiliano Trucchi

## Abstract

Insects are commonly expected to be protected from genomic erosion by high fecundity, short generation times, and large census sizes. Yet, insect populations can decline and eventually go extinct, underscoring the need for genomic indicators that provide actionable early warnings of population collapse. Here, we test this expectation by comparing contemporary and historical genomes of the Ponza grayling, *Hipparchia sbordonii*, an endangered butterfly endemic to the Pontine Islands in the Mediterranean, with the genomes of a widespread European congeneric species, *H. semele*. Using whole-genome resequencing, outgroup-based variant polarization, demographic reconstruction, runs of homozygosity, selection scans, and annotation-based genetic-load analyses, we show that *H. sbordonii* has undergone sustained demographic contraction, including a sharp recent decline. Despite limited temporal change in mean genome-wide heterozygosity, we observe extensive autozygosity, elevated inbreeding, and a clear shift from masked to realized genetic load in contemporary *H. sbordonii*. R’_XY_ analysis revealed similar relative frequencies of high-impact derived variants in *H. sbordonii* and *H. semele*, suggesting ineffective purging during population collapse, whereas low- and moderate-impact variants were relatively enriched in *H. sbordonii*, particularly within candidate regions under selection. This indicates a more complex dynamic, in which functional variation has been shaped by the combined effects of drift, relaxed purifying selection, and possible local adaptation. Our study shows that declining butterfly populations bear distinctive signatures of genomic erosion, mirroring patterns well documented in vertebrates. Yet recent demographic collapse in *H. sbordonii* is more clearly captured by long runs of homozygosity and realized genetic load than by changes in mean genome-wide heterozygosity, highlighting their potential as early warning indicators for monitoring declining insect populations.

## Introduction

Alarming declines in insect populations and biomass over the past three decades have raised concern about a potential ‘insect apocalypse’ (Hallmann et al. 2017; Lister & Garcia 2018; Sánchez-Bayo & Wyckhuys 2019; Forister et al. 2019; Cardoso & Leather 2019). Decreases in insect species are driven by a complex interplay of factors, including habitat loss, agricultural intensification, pesticide exposure, climate change, invasive species, and pollution, all of which impact communities at multiple scales (Webster et al. 2023; Wagner et al. 2021). As increasingly used as bioindicators to monitor ecosystem health (Chowdhury et al. 2023; Comay et al. 2021), butterflies are more clearly revealing their widespread population size collapse. In the United Kingdom, overall butterfly abundance has declined by about 50% since 1976. In the Netherlands and Flanders (Belgium), abundance has dropped by approximately 50% and 30%, respectively. Notably, a pan-European index shows a 39% decline in grassland butterfly populations since 1990 (Warren et al. 2021). Similarly, in the United States, total butterfly abundance declined by 22% between 2000 and 2020, with 13 times more species showing significant declines, and some species experiencing reductions exceeding 90% (Edwards et al. 2025). Despite their ecological importance, butterflies, and insects in general, are underrepresented in conservation efforts (Didham et al. 2020; Salvador et al. 2021; Dunn 2005), and often lack the suite of advanced genomics tools which are more commonly used in vertebrate conservation research (Webster et al. 2023; Sucháčková Bartoňová et al. 2023; Cardoso et al. 2011).

Understanding genetic diversity dynamics in declining populations is crucial for developing an early warning system to inform conservation strategies before irreversible genomic erosion occurs (Bosse & van Loon 2022; Díez-del-Molino et al. 2018). Declining population size can lead to a reduction in genetic diversity and an increase in inbreeding and genetic load (unmasking of recessive deleterious alleles), which compromise both population viability and adaptive potential (Bertorelle et al. 2022). As selection becomes less efficient at small effective population size, the interaction between genetic drift and purifying selection can determine whether deleterious alleles are purged or drift to higher frequency: while inbreeding may expose deleterious alleles to selection (purging), it may also lead to the fixation of harmful mutations, especially when effective population size is critically low (Dussex et al. 2023; Robinson et al. 2023).

Yet insects are rarely the object of conservation genomic studies (Webster et al. 2023), leaving it unclear whether they are experiencing the same patterns of genomic erosion documented in many vertebrate taxa. Unlike vertebrates, insect populations undergo strong demographic fluctuation and can remain relatively large, which could supposedly buffer them against the genomic consequences of small population size (Webster et al. 2023; Whitla et al. 2024). However, large or rapidly recovering census sizes may conceal temporary contractions into small local breeding groups. Under this scenario, increased mating among relatives during demographic contraction would promote autozygosity and the homozygous expression of derived functional variants, whereas residual or renewed gene flow during subsequent expansions could preserve population-wide diversity (Mattila et al. 2012; Ceballos et al. 2018; Dussex et al. 2023; Whitla et al. 2024; Nabholz et al. 2026). Strong demographic fluctuations and transient local fragmentation may therefore generate patterns of genomic erosion before conventional abundance estimates or mean genetic diversity metrics reveal a clear decline, emphasizing the need for sensitive genomic early-warning indicators capable of detecting cryptic genomic erosion before it becomes irreversible (Broquet et al. 2010; Jangjoo et al. 2020; Gompert et al. 2021; Hill et al. 2023; Whitla et al. 2024; Nabholz et al. 2026).

In this study, we investigate the genomic consequences of population decline in the Ponza grayling, *Hipparchia sbordonii* (Nymphalidae: Satyrinae), an endangered butterfly endemic to small Mediterranean islands. This species is restricted to the Pontine archipelago west of Naples, with a historical range limited to the three small islands of Ponza, Palmarola, and Zannone (Figure 1a), covering only about 16 km², but found only in Ponza in recent years. The main threat to its survival appears to be improper land use and biodiversity management (Bonelli et al. 2018). To investigate whether declining insect populations experience genomic erosion comparable to that documented in vertebrates, we compare the small, range-restricted population of *H. sbordonii* with its widespread congeneric *Hipparchia semele*, which maintains a much larger population size and occurs on the Italian mainland near the Pontine archipelago (Cesaroni et al. 1994). By analyzing both historical and contemporary samples, we assess which genomic metrics are most sensitive to the demographic collapse observed in the Ponza grayling, and whether they could serve more broadly as early warning signals for monitoring declining insect populations.

**Figure 1.**
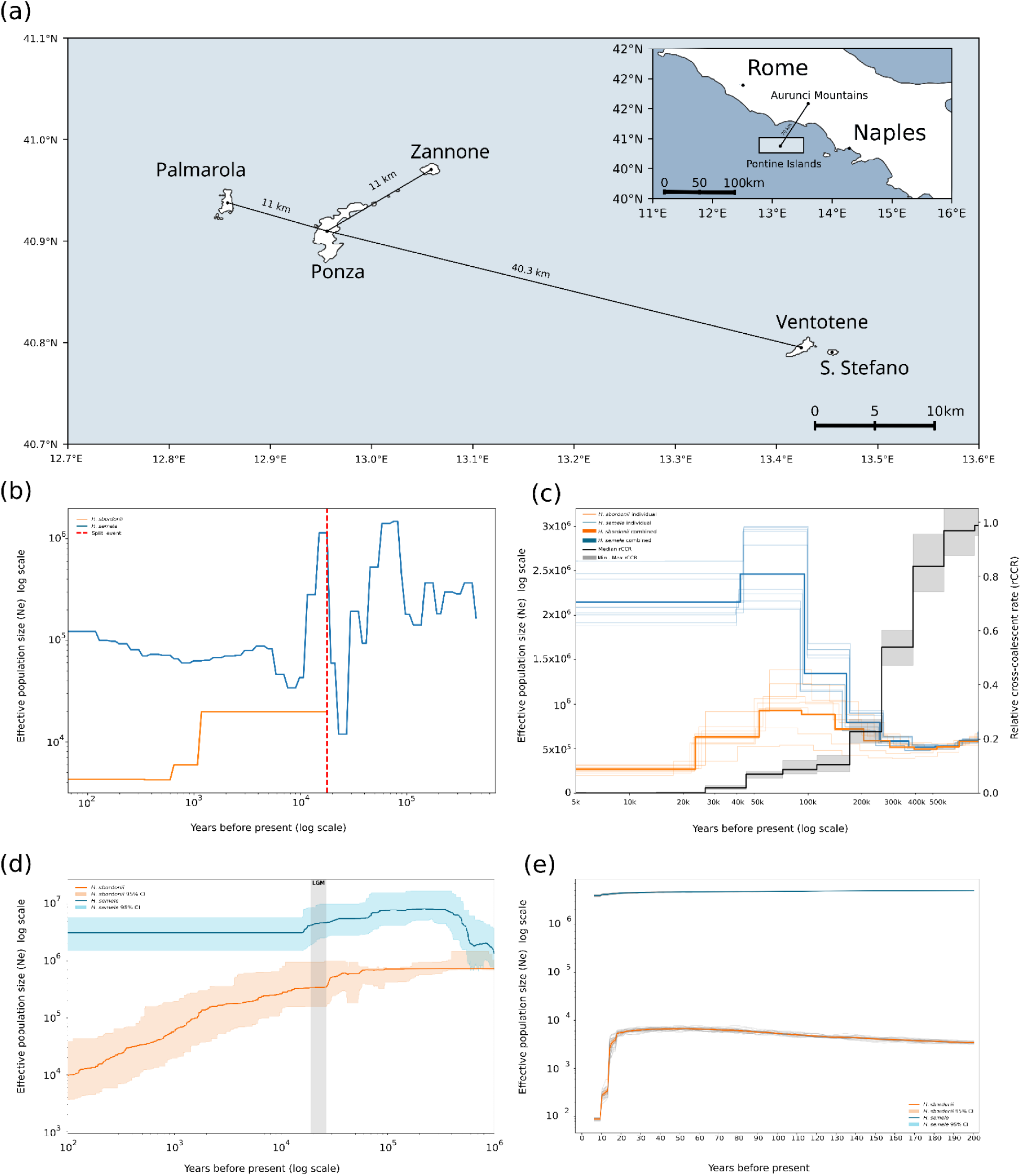
Demographic history of *H. sbordonii* and *H. semele* inferred using four complementary approaches. (a) Map of the Pontine Archipelago, showing the islands of Ponza, Palmarola, Zannone, Ventotene and Santo Stefano; the inset indicates the location of the archipelago along the central Italian coast, between the Aurunci Mountains and the Gulf of Naples. (b) *N*e trajectories reconstructed with SMC++, with the red dashed vertical line marking the estimated split between the two species. (c) MSMC2 estimates of effective population size (*N*e) through time for *H. sbordonii* (orange) and *H. semele* (blue). Thin lines represent individual-based trajectories, whereas bold lines indicate population-level estimates. The black line shows the median relative cross-coalescence rate (rCCR) between the two species, and the grey shaded area indicates the minimum-maximum range across cross-population runs. (d) *N*e trajectories reconstructed with Stairway Plot 2. Shaded areas indicate 95% confidence intervals based on 200 bootstrap replicates of the site frequency spectrum, and the grey vertical band marks the approximate timing of the Last Glacial Maximum (LGM) in Italy. (e) Recent changes in *N*e inferred with GONE from linkage disequilibrium patterns over the last approximately 200 generations. Dark lines indicate mean estimates and lighter lines the corresponding 95% confidence intervals.

## Results

### Long-term demographic contraction in *H. sbordonii* culminates in a sharp recent decline

Demographic history provides the baseline against which signatures of genomic erosion should be interpreted. Accordingly, we inferred and compared historical and recent effective population size trajectories in *H. sbordonii* and *H. semele* using Tajima’s D together with multiple complementary approaches (MSMC2, SMC++, Stairway Plot v2, and GONE). Tajima’s D analysis (Figure S1) indicated a slightly positive value in recent samples of *H. sbordonii* (median = 0.55), consistent with a population undergoing a recent bottleneck. In contrast, *H. semele* exhibited a strongly negative Tajima’s D (median = –1.34). Notably, in the historical samples of *H. sbordonii*, the median Tajima’s D was lower and close to zero, 0.22 for Palmarola and 0.26 for Ponza (Figure S1). Under a clean-split model, SMC++ estimates an effective split at ∼17,000 years ago (Figure 1b), likely during the final stages of the Last Glacial Maximum (Terhorst et al. 2017). The MSMC2 results (Figure 1c) provide the most detailed view of long-term population history. The effective population size (*Ne*) trajectories inferred from MSMC2 indicate a prolonged and continuous decline in *H. sbordonii*, beginning soon after divergence from *H. semele*, approximately 250,000 years ago. This pattern is consistently observed across individual replicates (Figure 1c), with *H. sbordonii* showing a steep reduction in *Ne* over time, reaching a low and stable plateau in recent history. In contrast, *H. semele* maintained relatively high and more fluctuating *Ne* values, with several expansion and contraction before stabilizing at a large effective population size. The MSMC2 relative cross-coalescence rate (rCCR) between *H. sbordonii* and *H. semele* showed a progressive decline, with rCCR dropping below 0.5 between ∼400,000 and 250,000 years ago and approaching zero around ∼15,000 years ago (Figure 1c), consistent with a gradual reduction in shared ancestry between *H. sbordonii* and *H. semele*. Together, these results support a scenario of progressive divergence culminating in a recent species split. Stairway Plot v2, parameterized with the folded site frequency spectrum computed for each species as input (Figure S2), similarly supports a long-term contraction in *H. sbordonii*, whereas *H. semele* retains a larger Ne with comparatively weaker fluctuations (Figure 1d). Although SMC++, Stairway Plot v2 and MSMC2 rely on different model assumptions and temporal resolutions, they show broadly concordant demographic trajectories. While the absolute effective population size (Ne) estimates differ across methods, they consistently identify a steady demographic decline in *H. sbordonii*, contrasted with a more variable but overall stable demography in *H. semele*. Notably, *H. sbordonii* shows no evidence of population recovery after the split, reinforcing its prolonged isolation and sustained demographic contraction. To investigate the most recent demographic trends, we used GONE. This analysis revealed a sharp reduction in effective population size in *H. sbordonii in* the last ∼30 generations (Figure 1e), further supporting the occurrence of a recent and severe bottleneck. In stark contrast, recent demographic inference for *H. semele* using GONE indicates no comparable decline: over the same timescale its effective population size has remained broadly stable, consistently hovering near 4 million, although GONE does detect a very recent downturn whose magnitude is much smaller than that inferred for *H. sbordonii*. Consistent with this recent demographic contraction, the temporal R_XY_ comparison between contemporary (2019) and historical (1990) Ponza samples showed higher derived allele frequencies in contemporary individuals for modifier, low-, and moderate-impact variants, whereas high-impact variants showed no significant temporal change (Figure S7). The increase observed also in putatively neutral modifier variants suggests that this pattern primarily reflects genome-wide demographic change associated with the recent decline.

### No evidence of gene flow from the mainland and low inter-islands connectivity in *H. sbordonii*

To investigate whether *H. sbordonii* and *H. semele* are genomically discrete lineages and whether *H. sbordonii* populations show genetic continuity through time, or evidence of recent connectivity among islands or with the mainland, we combined PCA, relatedness analyses, and pairwise F*_ST_*. PCA revealed a clear separation between *H. sbordonii* and *H. semele*, as well as between *H. sbordonii* samples from different islands and time points (Figure 2a). To avoid biasing the inference of population structure, we performed a relatedness analysis, and found that sample Ind_5 of *H. sbordonii* from Ponza (2019), was highly related to sample Ind_10 (pairwise relatedness value of 0.195; Figure S3). Thus, sample Ind_5 was excluded from the PCA. Within *H. sbordonii*, no major differentiation was observed between recent (2019) and historical (1990) samples from Ponza, consistent with the low F*_ST_* values. Pairwise F*_ST_* comparisons further confirmed genetic differentiation between the two species: the mean F*_ST_* between *H. sbordonii* from Ponza (2019) and *H. semele* was 0.186. Within *H. sbordonii*, comparisons between historical and recent samples revealed extremely low F*_ST_* values (e.g., F*_ST_* = 0.004 between Ponza 1990 and Ponza 2019). In contrast, slightly higher F*_ST_* values were observed between Ponza and Palmarola samples (e.g., F*_ST_* = 0.082 between Palmarola 1990 and Ponza 1990) (Figure S4).

**Figure 2.**
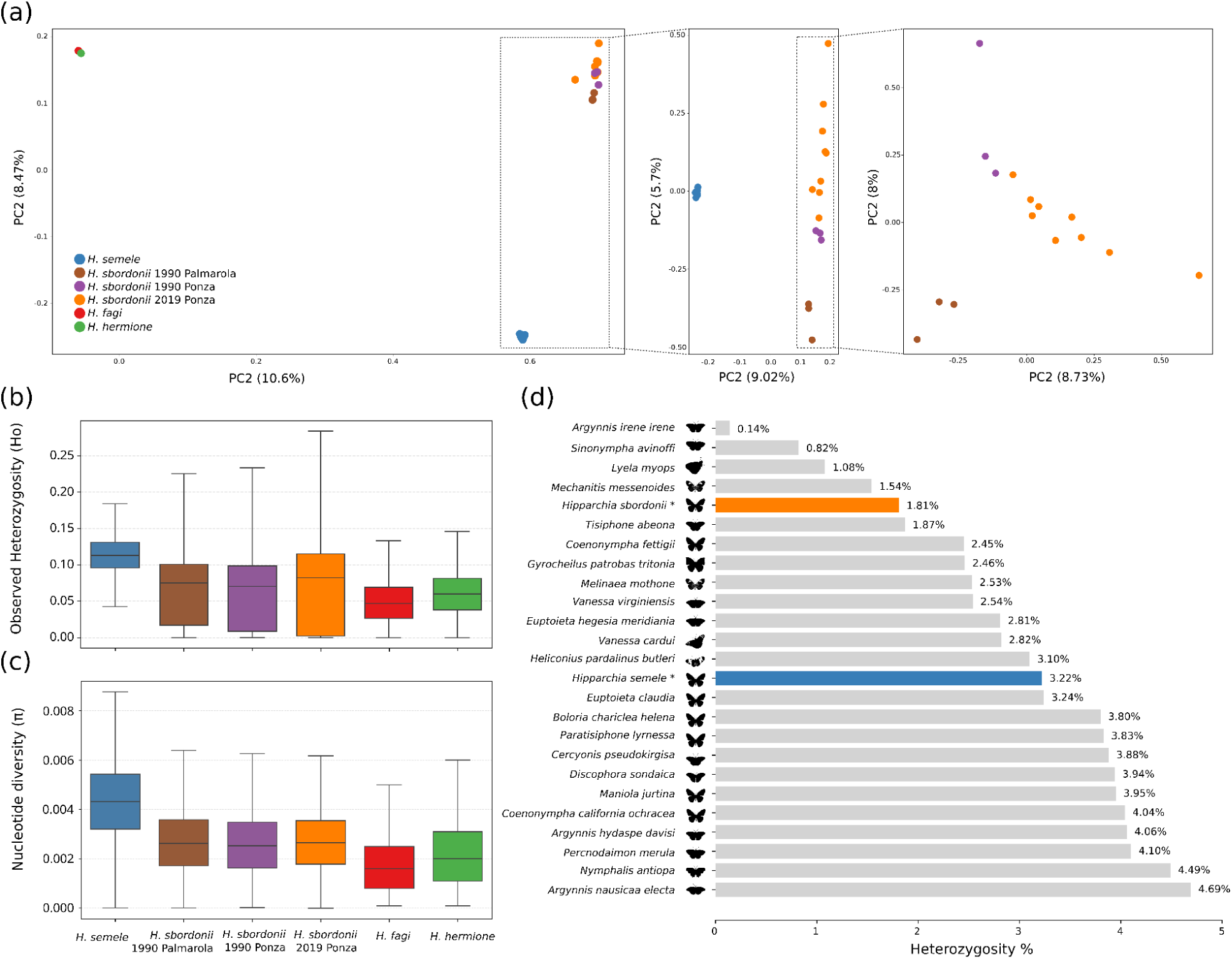
Genome-wide population structure, diversity, comparative heterozygosity in *Hipparchia* populations. (a) Principal Component Analyses (PCA) of genome-wide SNP variation after the exclusion of one highly related *H. sbordonii* individual from Ponza 2019. The three PCA plots show, from left to right: all samples including the outgroups *H. fagi* and *H. hermione*, only *H. sbordonii* and *H. semele*, and only *H. sbordonii* samples from Ponza 2019, Ponza 1990s, and Palmarola 1990s. (b) Observed heterozygosity (Ho) and (c) nucleotide diversity (π) estimated in non-overlapping 10 kb autosomal windows across *H. fagi*, *H. hermione*, *H. semele*, and three *H. sbordonii* populations. Boxplots show the median and interquartile range, with whiskers extending to 1.5×IQR. (d) Genome-wide heterozygosity estimates for *H. sbordonii*, *H. semele*, and *H. fagi* compared with publicly available estimates from other nymphalid butterflies. Asterisks indicate estimates generated from the present dataset.

### Reduced genetic diversity and elevated inbreeding in *H. sbordonii* relative to *H. semele*

To assess whether the demographic contraction inferred for the insular endemic *H. sbordonii* translated into genome-wide signatures of genomic erosion, we compared genetic diversity and inbreeding metrics with those of its widespread sister species *H. semele*. Specifically, we considered heterozygosity and nucleotide diversity (π) for genetic diversity and quantified recent inbreeding using runs of homozygosity (ROH) and F_ROH_. Despite the striking difference in recent effective population size between the two species, genetic diversity in *H. sbordonii* was only moderately reduced relative to *H. semele*. Recent *H. sbordonii* from Ponza showed approximately 38% lower observed heterozygosity and nucleotide diversity than *H. semele* (Figure 2b-c). Moreover, these values were comparable to, or higher than, those observed in the sampled *H. fagi* and *H. hermione* individuals. Specifically, the percentage of heterozygosity in *H. sbordonii* is lower compared to other Nymphalidae species (Figure 2d). Within *H. sbordonii*, however, recent samples from Ponza (2019) exhibited slightly higher median heterozygosity and π than historical samples from the same population collected in the 1990s, with historical values approximately 10% lower for heterozygosity and 4% lower for π. Despite this increase in central tendency, the 2019 genomes displayed more uneven heterozygosity landscapes, with a larger fraction of windows showing Ho = 0 and longer stretches of homozygosity, consistent with increased recent inbreeding (Figure 3, S5). Inbreeding levels were assessed by analyzing ROH across individuals from both *H. sbordonii* and *H. semele*. The results revealed that *H. sbordonii* individuals, particularly recent samples from Ponza, possess a substantially higher number and cumulative length of ROHs, especially in the longest size classes (>1 Mb), compared to *H. semele* (Figure 3). Notably, samples 5 and 10, which were previously found to be closely related, also exhibited extensive ROH. By contrast, ROHs were extremely rare and nearly absent in *H. semele*, resulting in very low cumulative ROH lengths and providing little evidence of recent inbreeding. To quantify inbreeding more directly, we calculated F_ROH_, the proportion of the genome in ROH across different ROH length thresholds (Figure 3). *H. sbordonii* exhibited notably higher F_ROH_ values across all ROH classes, with some individuals showing values consistent with recent parental relatedness. In contrast, *H. semele* displayed uniformly low F_ROH_ values, mostly confined to the shortest ROH class (<100 kb and 100–250 kb), further indicating little evidence of recent inbreeding (Supplementary Table 1). The independent PLINK analysis corroborated these patterns, yielding mean total FROH values of 0.191 for contemporary *H. sbordonii* from Ponza, compared with 0.139 and 0.111 for historical Ponza and Palmarola samples, respectively, and only 0.00056 for *H. semele*. ROHs longer than 1 Mb were detected exclusively in contemporary Ponza individuals, further supporting the marked increase in recent autozygosity inferred with BCFtools (Figure S6; Supplementary Table 2).

**Figure 3.**
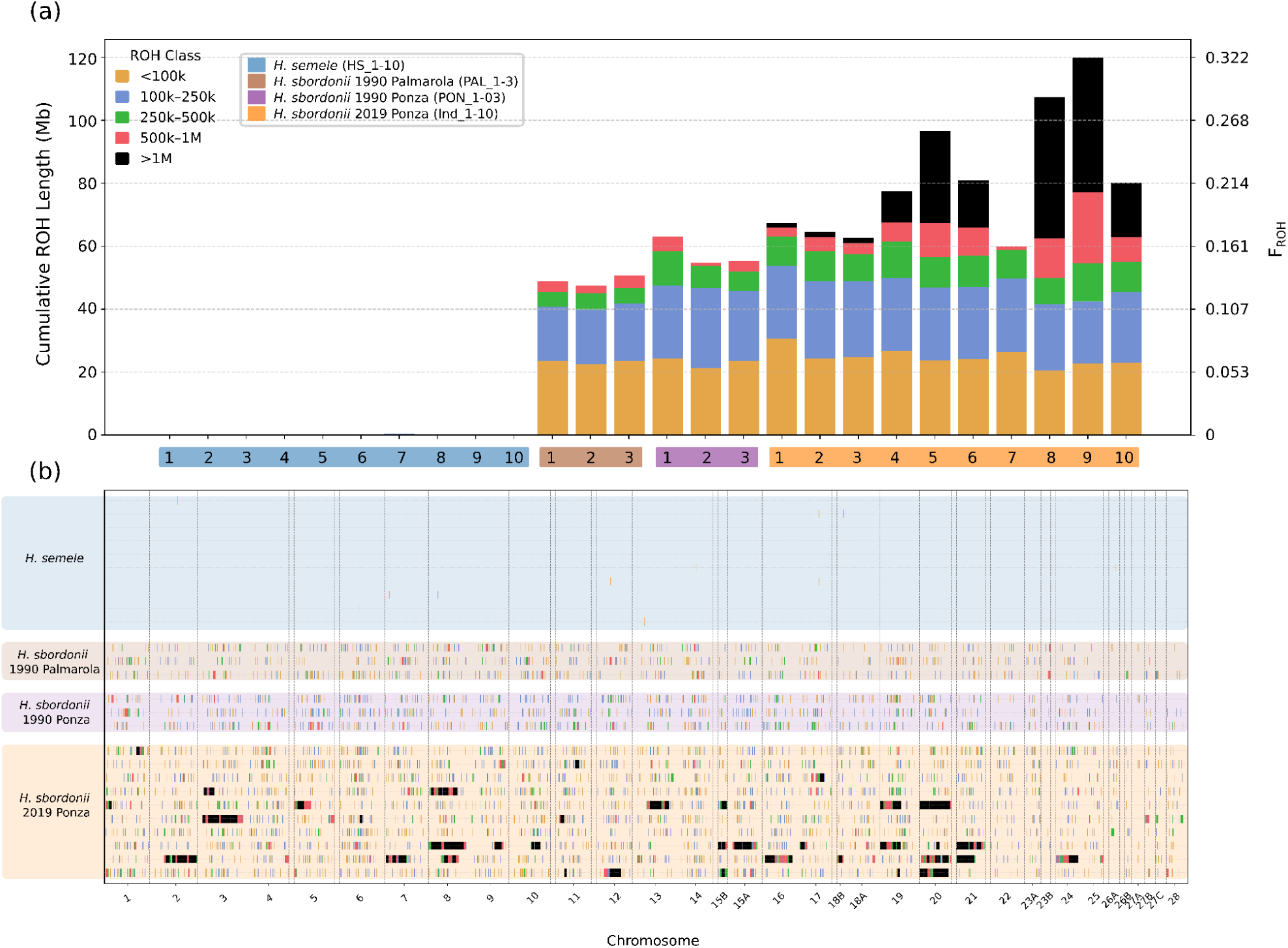
Genome-wide distribution of Runs of Homozygosity (ROH) and F_ROH_ across individuals and chromosomes. (a) Runs of homozygosity (ROH) detected from high-quality autosomal SNPs using BCFtools roh. Stacked bars show the cumulative ROH length for each individual, partitioned into five length classes: <100 kb, 100–250 kb, 250–500 kb, 500 kb–1 Mb, and >1 Mb. The secondary y-axis reports F_ROH_, calculated as the proportion of the autosomal genome contained within ROH. Individuals are grouped by population and sampling period: mainland *H. semele* (2019), contemporary *H. sbordonii* from Ponza (2019), and historical *H. sbordonii* from Ponza and Palmarola (1990). (b) Genome-wide distribution of ROH across autosomal chromosomes for the same individuals. Each horizontal track represents one individual, and colored segments indicate ROH according to the same five length classes shown in panel (a). Vertical dashed lines delimit chromosomes.

### Inbreeding shifts derived functional variation from masked to realized load without detectable purging

We tested whether the recent contraction and elevated inbreeding in *H. sbordonii* translate into a shift in the balance between realized and masked load by quantifying homozygous and heterozygous derived deleterious variation and estimating the R_XY_ index across different fitness effect categories. The comparison of realized and masked genetic load between *H. sbordonii* and *H. semele* highlights clear differences in how deleterious mutations are distributed. *H. sbordonii*, which has undergone a recent demographic decline and has elevated inbreeding, shows a higher realized load, with a greater number of homozygous derived variants (Figure 4b). Conversely, *H. semele*, with its larger and more stable population, exhibits a higher masked load, with more deleterious alleles remaining in a heterozygous state (Figure 4a). To investigate the evolutionary dynamics of deleterious alleles between the two species, we applied both the R_XY_ and R′_XY_ statistics. R_XY_ measures the ratio of derived allele frequencies between two populations across categories of mutational impact, with X = *H. sbordonii* and Y = *H. semele*. In our case, we observed R_XY_ < 1 from modifier (putatively neutral) to High-effect mutations (Figure 4e), indicating that *H. sbordonii* has lower frequencies of derived alleles than *H. semele* across all categories. When we examined R_XY_ values separately for homozygous and heterozygous genotypes, we observed that R_XY_hom values remained above 1 across all categories, while R_XY_het values were consistently lower in *H. sbordonii* (Figure 4c–d). This pattern reflects the loss of heterozygosity in *H. sbordonii* due to its demographic history, rather than a true reduction in overall derived allele content.

**Figure 4.**
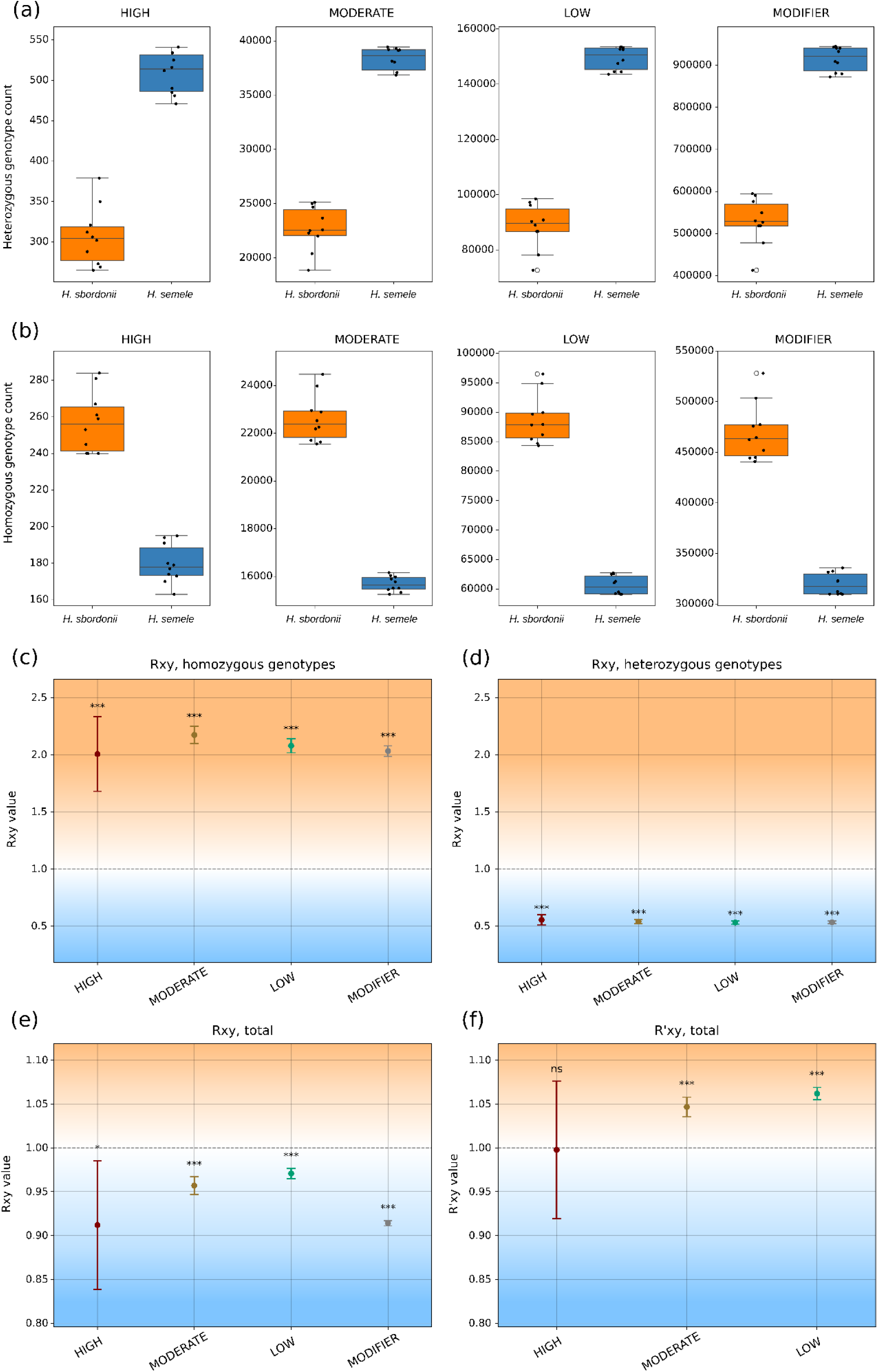
Genotype-based genetic load and R_XY_/R′_XY_ comparisons between *H. sbordonii* and *H. semele*. (a) Per-individual counts of heterozygous genotypes and (b) homozygous derived genotypes across four mutational effect categories: high, moderate, low and modifier. Boxplots summarize variation among individuals, with points representing single samples.(c-f) R_XY_ and R’_XY_ estimates are based on a weighted leave-one-block-out jackknife across 100 genomically ordered blocks. (c) R_XY_ values calculated from homozygous derived genotypes, used as a proxy for relative realized load. (d) R_XY_ values calculated from heterozygous genotypes, used as a proxy for relative masked load. (e) Total R_XY_ values calculated from derived allele frequencies across mutational effect categories. (f) R′_XY_ values calculated from derived allele frequencies and normalized by putatively neutral modifier variants to account for genome-wide demographic differences between populations. Variants were annotated with SnpEff and grouped according to predicted functional impact. SNPs were polarized using *H. fagi* and *H. hermione* as outgroups to infer ancestral and derived states. R_XY_ statistics were computed following Do et al. (2015), and uncertainty was estimated using a weighted 100-block jackknife approach. Points in panels c-f indicate weighted jackknife estimates, while error bars represent 95% confidence intervals. The dashed horizontal line indicates the neutral expectation of R_XY_ or R′_XY_ = 1. Asterisks indicate significant deviations from 1 after FDR correction (*** adjusted p < 0.001), while “ns” indicates non-significant differences. Two-tailed P values for deviations from the null expectation of R_XY_ or R′_XY_ = 1 were calculated from the weighted block-jackknife estimates and standard errors using a t distribution with 99 degrees of freedom and adjusted for multiple testing using the Benjamini–Hochberg procedure.

To test for signatures of purging relative to the large mainland population of *H. semele*, we estimated the normalized R’_XY_ index which can highlight different accumulation of non-neutral variants relative to neutral ones. R′_XY_ estimate for high-effect variants was close to 1 and showed no evidence of depletion in *H. sbordonii* relative to *H. semele* (weighted block-jackknife estimate = 0.998, SE = 0.039, 95% CI: 0.919–1.076; Figure 4f). Thus, although increased homozygosity in small inbred populations can expose recessive deleterious mutations to purifying selection, our data do not support detectable purging of strongly deleterious variation in *H. sbordonii*. If any depletion of high-effect variants exists, it is too small relative to the uncertainty of the estimate to be distinguished from the demographic baseline (e.g. Grossen et al. 2020). The temporal comparison between contemporary and historical Ponza samples similarly showed no significant deviation of R’_XY_ from 1 for any functional class, providing no evidence of recent purging (Figure S7). By contrast, in the interspecific comparison between *H. sbordonii* and *H. semele*, moderate- and low-effect mutations showed R′_XY_ values significantly greater than 1. The increase was clear for moderate-effect variants (weighted block-jackknife estimate = 1.047, SE = 0.006, 95% CI: 1.036–1.058; Figure 4f) and even stronger for low-effect variants (weighted block-jackknife estimate = 1.062, SE = 0.003, 95% CI: 1.055–1.069; Figure 4f). Thus, after correction using putatively neutral modifier variants, *H. sbordonii* retains a relative excess of low- and moderate-effect derived mutations compared with *H. semele*.

### Candidate regions under selection are associated with elevated R′_XY_ in low and moderate functional variants

To assess whether the excess of low- and moderate-effect derived variants in *H. sbordonii* was associated with genomic regions showing signatures of selection, we intersected polarized functional SNPs with high-confidence candidate regions identified by integrating CLR, iHS, positive XP-EHH, Tajima’s D, reduced nucleotide diversity, and elevated F*_ST_*. This combined scan identified 34 putative selection candidate regions, spanning approximately 2.43 Mb across 17 chromosomes (Figure 5). Twelve regions were supported by all six metrics, whereas the remaining 22 were supported by five metrics (Figure S8). All retained regions included support from at least five of the six metrics. The 34 regions under selection overlapped 10,140 functional SNPs, all located in genic regions and mapping to 168 genes. Most of these SNPs were classified as moderate- or low-impact variants, whereas only a smaller fraction corresponded to high-impact variants. We then tested whether R′_XY_ values for low- and moderate-effect derived variants were higher within these regions under selection than in the genomic background. The bootstrap analysis revealed consistently higher R′_XY_ values inside regions under selection for both functional categories (Figure 6b). This increase was more pronounced for moderate-effect variants, whereas low-effect variants showed a smaller but still significant increase (Figure 6b).

**Figure 5.**
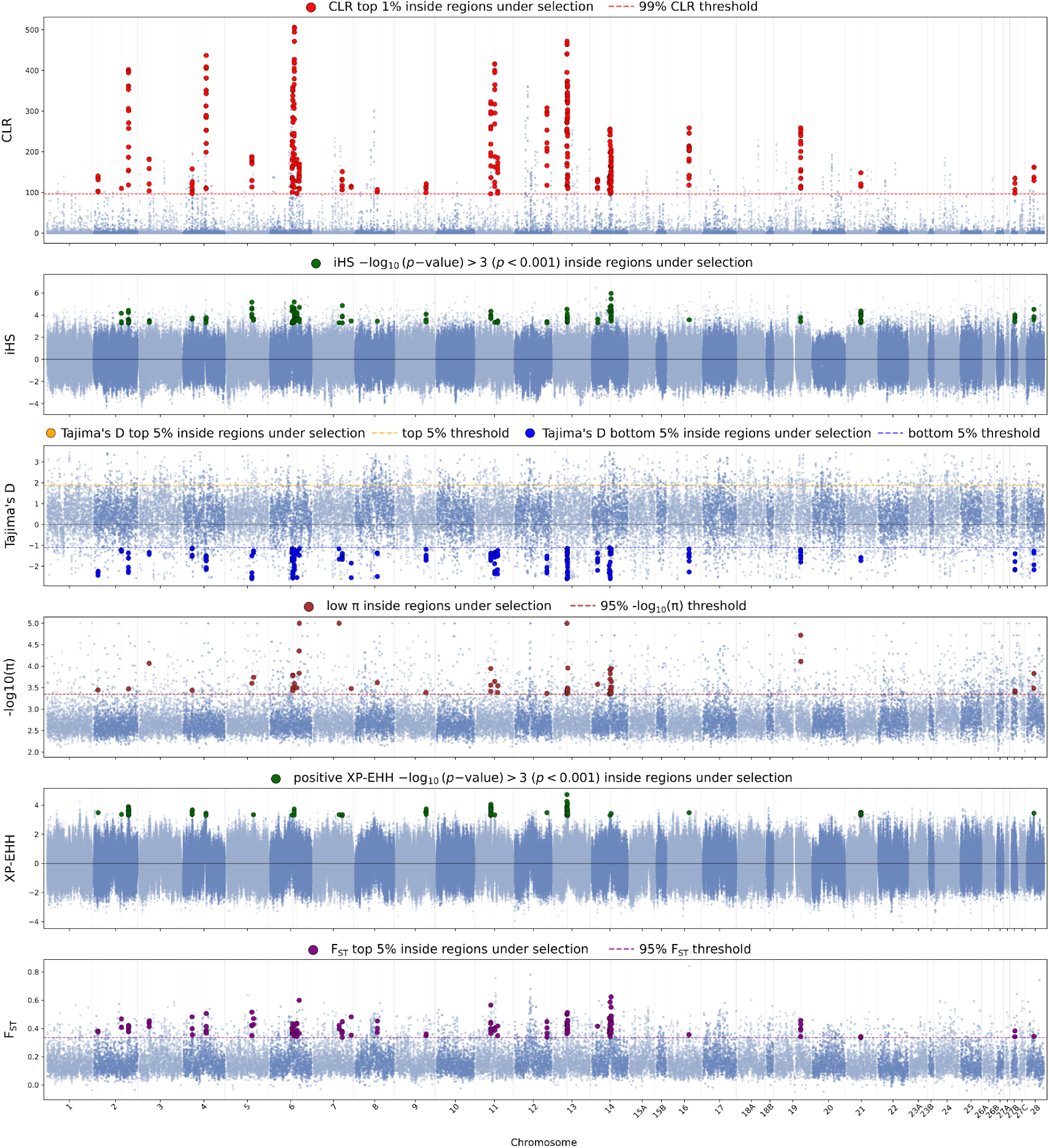
Genome-wide distribution of selection and differentiation statistics used to identify high-confidence candidate regions in *Hipparchia sbordonii*. Manhattan-style plots showing, from top to bottom, CLR from SweepFinder2, iHS, Tajima’s D, -log_10_(π), positive XP-EHH and F*_ST_* across autosomal chromosomes. Pale blue points represent genome-wide values, while coloured points indicate threshold-passing values located within high-confidence candidate regions. Thresholds are shown by dashed horizontal lines for CLR, Tajima’s D, -log_10_(π) and F*_ST_*. Candidate signals were defined as the top 1% of CLR values, iHS sites with -log10(p-value) > 3, the lower and upper 5% tails of Tajima’s D, the upper 5% of -log_10_(π), positive XP-EHH sites with -log_10_(p-value) > 3, and the upper 5% of F*_ST_*. These regions show concordant signals of reduced diversity, negative Tajima’s D, elevated differentiation and extended haplotype homozygosity, consistent with candidate loci under recent or ongoing selection in *H. sbordonii*.

**Figure 6.**
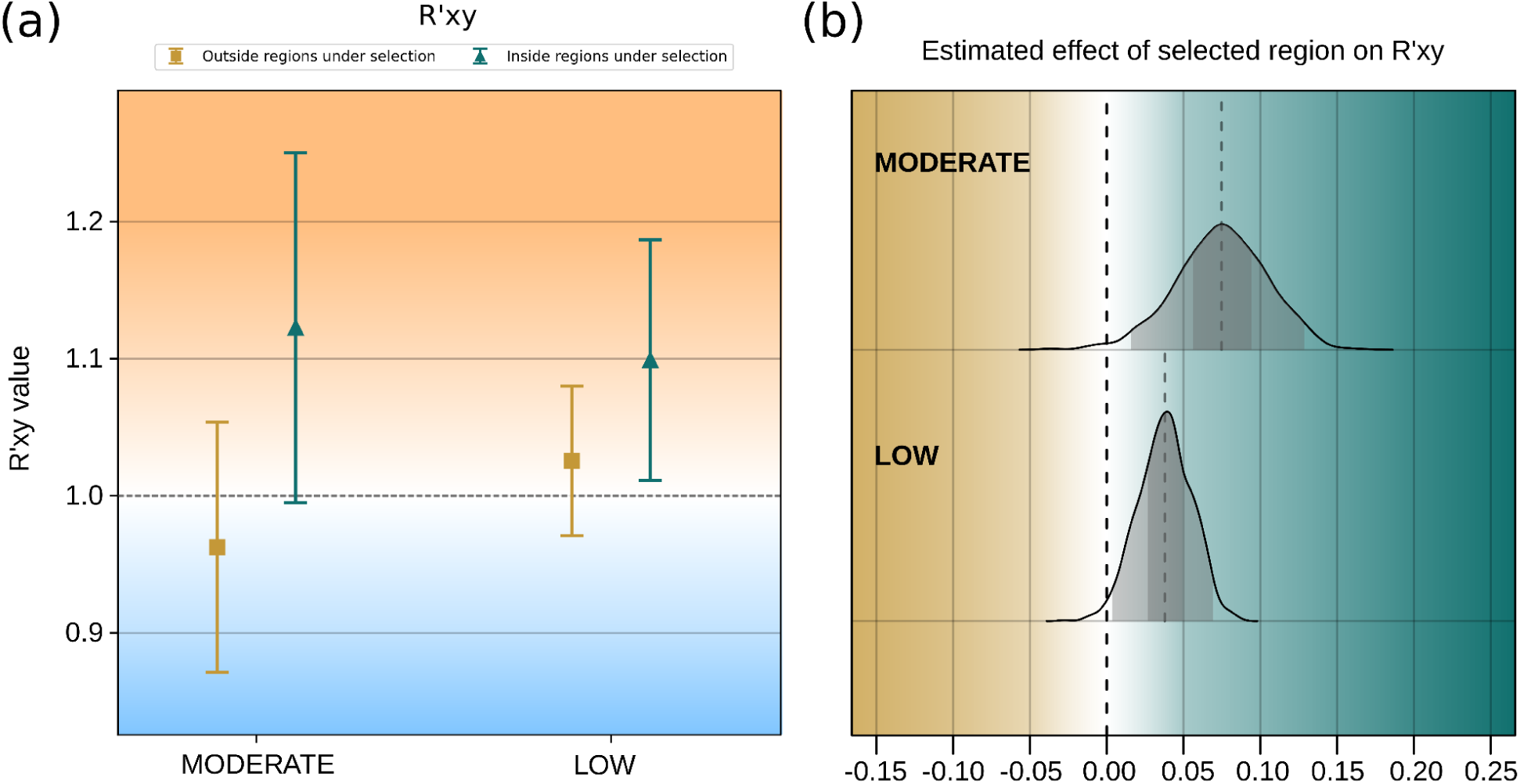
Effect-matched comparison of R′_XY_ estimates inside and outside candidate regions under selection. (a) Representative single resampling iteration showing R′_XY_ estimates for SNPs located inside candidate regions under selection and for an effect-matched genomic background outside these regions. For visualization, outside-region SNPs were randomly subsampled to match the number of SNPs observed inside candidate regions for each functional category, preserving the relative functional composition of the comparison. R′_XY_ was obtained by normalizing R_XY_ values by the corresponding modifier class. Points represent weighted block-jackknife estimates across 100 genomically ordered blocks, and error bars indicate 95% confidence intervals. The dashed horizontal line indicates the neutral expectation of R′_XY_ = 1. Only moderate- and low-effect variants are shown. (b) Distribution of linear model coefficients estimated from the full bootstrap analysis used to test differences in R′_XY_ inside versus outside candidate regions under selection. In this analysis, outside-region SNPs were sampled at three times the number of SNPs observed inside candidate regions for each functional category, as described in the Methods. Density curves represent bootstrap distributions of effect estimates across 1,000 iterations for low- and moderate-effect variants. Shaded areas indicate the central 50% and 95% of the bootstrap distribution.

BlastKOALA assigned KEGG Orthology (KO) terms to 100 of the 168 genes, 56 of which were associated with at least one KEGG pathway. KOBAS enrichment analysis identified significant terms related to lipid transport, ecdysone biosynthesis, chitin-based cuticle formation, ATPase-coupled transmembrane transport, and signalling receptor activity, while KEGG and Reactome results also implicated ABC transporters and COPII-mediated vesicle transport (Figure S9). Inspection of the BlastKOALA KO/KEGG annotations further highlighted genes involved in membrane transport, hormone biosynthesis, mitochondrial metabolism, proteostasis, DNA/RNA maintenance, and neuronal or sensory functions, including DIB/CYP302A1, ABCA3-like transporters, glutamate receptor-related genes, proteasome components, DNA replication factors, and mitochondrial respiratory genes.

## Discussion

While genomic erosion is well documented in vertebrates, insects remain underrepresented in conservation genomics, particularly endemic, range-restricted, and declining taxa. Their high fecundity, short generation times, and often large census sizes have been proposed to buffer them from the detrimental effects of drift and inbreeding (Webster et al. 2023; Leung et al. 2025). However, insect populations can also undergo strong and rapid fluctuations in abundance, with recurrent contractions, fragmentation, and subsequent demographic recovery altering the balance between drift, inbreeding, and gene flow over relatively short timescales (Jangjoo et al. 2020; Gompert et al. 2021; Hill et al. 2023). Such dynamics may allow census abundance or mean genomic diversity to remain relatively high while local effective population size and mating patterns deteriorate. Our results show that signatures of a more subtle genomic erosion are appearing in *H. sbordonii*. Long ROH, elevated F_ROH_, and increased homozygosity of derived functional variants provided clearer evidence of recent genomic deterioration than mean genome-wide diversity, consistent with a similar decoupling between diversity and recent inbreeding reported in other range-restricted butterflies (Whitla et al. 2024; Nabholz et al. 2026). ROH-based autozygosity and relative genetic load may therefore provide particularly sensitive early-warning signals of genomic deterioration in insects whose strongly fluctuating census sizes can obscure recent decline in genome-wide diversity.

### Runs of homozygosity capture recent inbreeding more clearly than genome-wide diversity

The diversity results are consistent with the demographic history, but also highlight why genome-wide heterozygosity and nucleotide diversity should not be interpreted in isolation from ROH-based metrics (McQuillan et al. 2008; Kardos et al. 2015; Ceballos et al. 2018). Despite the striking difference in recent effective population size, *H. sbordonii* showed only moderately lower heterozygosity and nucleotide diversity than *H. semele* and ranked among the least genetically diverse Nymphalidae examined. However, among the *Hipparchia* samples analyzed, similarly low or even lower diversity values were observed in the sampled *H. fagi* and *H. hermione* individuals, both belonging to species classified as Least Concern in the Italian IUCN Red List. Moreover, the comparison between historical and contemporary Ponza samples revealed no pronounced temporal erosion of genome-wide diversity, with contemporary individuals showing instead slightly higher heterozygosity and π than those sampled in the 1990s, despite carrying a substantially greater burden of long ROH. This apparent discrepancy arises because ROH-based metrics capture autozygosity and recent shared ancestry, whereas mean heterozygosity and π summarize diversity across the whole genome and can remain relatively high outside homozygous tracts. Accordingly, long ROHs provide a sensitive signal of recent inbreeding and acute demographic contraction (Purfield et al. 2012; Ceballos et al. 2018). This pattern was particularly evident in some individuals that were also highly related and shared extensive ROHs, consistent with recent parental relatedness and potentially limited mate choice in a declining population (Kardos et al. 2015; Ceballos et al. 2018; Ringbauer et al. 2021). Overall, F_ROH_ provides a direct genome-wide estimate of realized autozygosity and recent inbreeding that complements genome-wide diversity summaries (McQuillan et al. 2008; Kardos et al. 2015; Ceballos et al. 2018). Beyond this difference in what the metrics capture, a complementary demographic mechanism may explain how high autozygosity can coexist with relatively preserved genomic diversity in insects. Recurrent demographic fluctuations and episodic fragmentation into small local breeding groups may reduce local population size and strengthen drift during sub-population contractions (Broquet et al. 2010; Jangjoo et al. 2016; Hill et al. 2023). Restricted dispersal and mating among relatives may then increase autozygosity and generate long ROH, while recessive deleterious variants in homozygous state can be exposed (and likely purged) in some but not all of the sub-populations due to the randomness of the genetic drift (Mattila et al. 2012; Ceballos et al. 2018; Dussex et al. 2023; Whitla et al. 2024; Nabholz et al. 2026). During subsequent recovery, restored connectivity and renewed gene flow may preserve or recover population-wide diversity (Jangjoo et al. 2016; Jangjoo et al. 2020; Gompert et al. 2021). Strong census fluctuations may therefore produce substantial recent autozygosity without an equally pronounced decline in mean heterozygosity, making ROH-based metrics a potentially more sensitive warning signal of recent genomic erosion (Webster et al. 2023; Whitla et al. 2024; Nabholz et al. 2026).

### Inbreeding shifts functional variation from masked to realized load without detectable purging

The assessment of genetic load shows the expected shift from masked to realized load under increasing homozygosity in the declining *H. sbordonii*. *H. semele*, which retains a much larger effective population size, carries more derived functional variants in the heterozygous state, consistent with a higher masked load. In contrast, *H. sbordonii* carries more homozygous derived functional variants, consistent with the exposure of recessive deleterious alleles during inbreeding (Figure 4a-b) (Bertorelle et al. 2022; Dussex et al. 2023). The global R_XY_ values below 1 at first glance might suggest fewer derived variants in *H. sbordonii*, but the genotype-specific analysis shows that this pattern is largely driven by the loss of heterozygosity (Figure 4c-e). Since bottlenecks, drift, and inbreeding alter the frequency of derived variants across the genome, including at putatively neutral sites, normalizing functional categories by putatively neutral modifier variants provides a demographic baseline against which deviations in functional derived variation can be interpreted. Thus, without this correction, R_XY_ would confound selection with demographic differences between the two species (Do et al. 2015; Xue et al. 2015; Grossen et al. 2020).

After this correction, the R′_XY_ results provide no evidence for a depletion of high-effect derived mutations in *H. sbordonii* (Figure 4f). Although increased homozygosity in small inbred populations can expose recessive deleterious alleles to purifying selection, our results do not support detectable purging of strongly deleterious variation in *H. sbordonii*. This interpretation is supported by the temporal comparison between historical and contemporary Ponza samples: although R_XY_ increased for moderate-, low-effect, and modifier variants, R’_XY_ did not differ from 1 for any functional class (Supplementary Figure 6), suggesting that these changes largely reflect genome-wide demographic effects and increased drift rather than purging. This temporal comparison should nevertheless be interpreted cautiously because it contrasts 10 contemporary individuals with only three historical samples.

In the interspecific comparison, if any depletion of high-effect variants exists, it is too small relative to the uncertainty of the estimate to be distinguished from the demographic background. The absence of a detectable depletion signal may reflect both the recent timing and the fluctuating nature of the demographic contraction. Effective population size may not have remained sufficiently low for enough generations to produce sustained inbreeding and repeated homozygous exposure of strongly recessive deleterious alleles, which are required for purging to generate a detectable genome-wide signal (Grossen et al. 2020; Khan et al. 2021; Dussex et al. 2023). Moreover, across fragmented populations subjected to independent genetic drift, it is less likely that a deleterious allele will reach high frequency, and then be more prone to purging, in all of the sub-population, thus increasing its chances of randomly surviving until demographic recovery or renewed connectivity. A second, non-mutually exclusive explanation comes from the exploratory analysis of the level of RNA expression per variant reported in the Supplementary Material (Figure S9). High-impact variants tend to occur in low-expression genomic contexts regions in both species, consistent with evidence that gene expression strengthens purifying selection and reduces the segregation of deleterious variants in highly expressed genes (Trucchi et al. 2025). Thus, although these variants are annotated as high-impact, the subset that remains segregating (and therefore not purged) may experience a weaker component of purifying selection due to their low expression rate. Under the stronger drift associated with the recent contraction of *H. sbordonii*, such low-expression high-impact variants may persist and rise in frequency, helping explain why R′_XY_ is close to 1 rather than showing a clear depletion.

### Low- and moderate-impact variation reflects both genomic erosion and adaptive differentiation

In contrast to high-impact variants, R′_XY_ values were significantly greater than 1 for low- and moderate-impact variants, indicating an excess of these functional categories in *H. sbordonii* when correcting for population-specific demography. This excess was stronger for low-effect variants and also evident for moderate-effect variants, suggesting that the clearest signal is not a generalized depletion of derived functional variation, but rather an accumulation of weaker-effect derived mutations in the declining *H. sbordonii* population. This pattern is consistent with theoretical expectations and empirical examples in which severe reductions in effective population size can reduce the efficacy of purifying selection, allowing mildly deleterious variants to drift to higher frequency (Bertorelle et al. 2022; Dussex et al. 2023).

However, the selection analyses, coupled with the R′_XY_ bootstrap comparison inside and outside regions under selection, provide a secondary layer of interpretation to the genetic load results. The consistent increase in R′_XY_ within regions under selection for both low- and moderate-impact variants suggests that part of the functional variation enriched in *H. sbordonii* is concentrated in genomic regions showing signals of selection across multiple tests (Figure 6). As a consequence, the excess of moderate-impact variants in *H. sbordonii* should not always be interpreted as genetic load alone. SnpEff impact categories describe predicted molecular consequences, but they do not directly define the fitness effect of each variant. Low- and moderate-impact variants may therefore include neutral, weakly deleterious, or potentially adaptive variation.

Thus, the excess of low- and moderate-impact variants in *H. sbordonii* may reflect a combination of relaxed purifying selection and genetic drift, but also adaptive differentiation. The enrichment observed within regions under selection suggests that at least part of the signal may be associated with local adaptation, either through direct selection on functional variants or through linked hitchhiking of nearby variants (Chun & Fay 2011; Marsden et al. 2016; Wang et al. 2024)

Functional annotations provide biological plausibility for a partial contribution of selection to the observed excess of low- and moderate-impact variants. Genes located in regions under selection were associated with membrane transport, ecdysone-related development, chitin-based cuticle formation, sensory or neuronal functions, mitochondrial metabolism, proteostasis, and DNA/RNA maintenance. These functions may be relevant in an insular context, where microclimate, host-plant use, phenology, dispersal constraints, and ecological interactions can differ from mainland environments. In particular, *DIB/CYP302A1*, involved in insect ecdysteroid biosynthesis and developmental regulation, and ABC transporter-like genes, which have broad roles in substrate transport and physiological responses in arthropods, represent plausible candidates contributing to local adaptation in *H. sbordonii* (Chávez et al., 2000; Dermauw & Van Leeuwen, 2014; Niwa & Niwa, 2014).

### Implications for insect conservation genomics

More broadly, our results address whether declining insects undergo genomic processes comparable to those documented in vertebrates, and which genomic metrics are most responsive to population decline. *H. sbordonii* shows that demographic contraction in insects can be associated with reduced diversity, increased inbreeding, extensive ROH, and altered patterns of derived functional variation (Webster et al. 2023; Beaurepaire et al. 2024; Leung et al. 2025). Similar patterns have recently been reported in other declining, isolated, and range-restricted butterflies (Whitla et al. 2024; De-Dios et al. 2024; Nolen et al. 2024; Nabholz et al. 2026). However, genomic outcomes are not uniform. They may depend on the severity, duration, and timing of demographic change, as well as on the degree of isolation and connectivity among populations (Webster et al. 2023; Macdonald et al. 2025). Among the metrics examined here, long ROH and high F_ROH_ provided the clearest evidence of recent genomic deterioration. Mean heterozygosity and π showed no pronounced temporal decline, despite substantially greater autozygosity in contemporary individuals (Kardos et al., 2015; Ceballos et al. 2018). Strong census fluctuations and episodic fragmentation may further increase this mismatch. Inbreeding may rise during population contractions, while renewed gene flow during recovery preserves population-wide diversity (Broquet et al. 2010; Jangjoo et al. 2016; Jangjoo et al. 2020; Gompert et al. 2021; Hill et al. 2023).

Functional variation provided a complementary view of genomic deterioration. In *H. sbordonii*, the shift from heterozygous to homozygous derived variants indicates that inbreeding is exposing previously masked functional variation. The absence of depletion among high-impact variants may reflect a contraction that was too recent, fluctuating, or insufficiently severe and sustained to generate repeated homozygous exposure and detectable purging (Grossen et al. 2020; Khan et al. 2021; Dussex et al. 2023; Jangjoo et al. 2016). Moreover, the segregating high-impact variants tended to occur in lowly expressed genomic contexts, where expression-associated purifying selection may be weaker (Trucchi et al. 2025), although this result remains exploratory. The excess of low- and moderate-impact variants may partly reflect drift and relaxed purifying selection, but their stronger signal within candidate regions under selection suggests that direct or linked selection may also contribute (Bertorelle et al. 2022; Dussex et al. 2023; Chun & Fay 2011; Marsden et al. 2016; Wang et al. 2024). ROH-based FROH and realized load may therefore provide valuable early-warning indicators of genomic deterioration in insects, particularly when census abundance fluctuates strongly and mean genome-wide diversity shows limited temporal change (Webster et al. 2023; Whitla et al. 2024; Nabholz et al. 2026). While conservation actions should maintain habitat quality, prevent further population fragmentation, preserve or restore connectivity among breeding groups, and avoid additional reductions in effective population size in *H. sbordonii*, genomic monitoring should be routinely integrated with field surveys and abundance estimates in every butterfly or insect-focused conservation study.

## Materials and Methods

### 1 Population sampling and genomics data generation

A total of 28 individuals were sampled for genomic analyses, including ten and three *H. sbordonii* specimens sampled in Ponza in 2019 (n=10: Ind_1-10) and 1990 (n=3: PON_1-3), respectively, three from Palmarola 1990 (n=3: PAL_1-3), and ten *H. semele* (n=10: HS_1-10) from the Italian mainland (Aurunci Mountains, Lazio, 2019) (Figure 1a). Two additional outgroup species, *H. fagi* (n=1) and *H. hermione* (n=1) (Valle Cupola, Lazio, 2019) were also included for comparative analyses and to assess the correct ancestral state of each SNP (polarization).

Contemporary butterfly samples were collected using hand nets and flash frozen in liquid nitrogen in the field. Historical samples from Ponza and Palmarola have been preserved frozen (−40°C) since their collection in 1990 at Tor Vergata University.

Whole DNA was extracted using a PureLink Genomic DNA Mini Kit (Invitrogen). DNA integrity was assessed by 1.5% agarose gel electrophoresis, and DNA concentration was measured using a Qubit 4 fluorometer Broad Range Assay (Invitrogen). Genomic libraries were constructed using a DNA PCR-Free Prep Kit (Illumina) according to the manufacturer’s protocol. Target coverage was 10-15X for all samples. Libraries were sequenced in paired-end mode on an Illumina NovaSeq 6000 System using a 300-cycle S2 Reagent Kit v1.5, generating 150 bp reads.

### 2 Read mapping, variant calling and filtering

All resequencing data were aligned to the chromosome-level *H. sbordonii* reference genome (Fava et al. 2024) using bwa-mem2 (v2.2.1) (Vasimuddin et al. 2019) with default parameters. Resulting alignments were sorted and indexed using samtools (v1.17) (Danecek et al. 2021). To ensure file integrity prior to downstream processing, samtools quickcheck was used on the resulting BAM files. Read groups were then assigned using bamaddrg, and optical and sequencing duplicates were marked using the MarkDuplicates module from Picard Tools (v3.2.0) (https://github.com/broadinstitute/picard). After the mapping, we obtained an average sequencing depth of 17X for *H. semele*, 14.5X for *H. sbordonii* (18X for recent individuals and 8.8X for historical individuals), 14.3X for *H. fagi*, and 12X for *H. hermione*, ensuring high-quality SNP calls. Variant calling was conducted using GATK v4.2.2.0 (McKenna et al. 2010) following the best practices pipeline for short variant discovery. The pipeline consists of generating per-sample GVCF files with HaplotypeCaller (-ERC GVCF mode), consolidating them using GenomicsDBImport to improve scalability, and performing joint genotyping with GenotypeGVCFs to obtain a raw multi-sample VCF. From the raw VCF, SNPs and indels were extracted separately using GATK SelectVariants and filtered in subsequent steps. Indels were filtered using VariantFiltration. SNPs were filtered using the same tool with the following criteria: QUAL < 60, QD < 2.0, FS > 60.0, MQ < 40.0, MQRankSum < −20.0, ReadPosRankSum < −8.0, and GQ < 10. SNPs located within 5 bp of high-confidence indels were masked using the --mask option in VariantFiltration. Repeated regions were masked using VCFtools (Danecek et al. 2011) and a custom fasta-like mask file, and all SNPs falling within those regions were excluded. After filtering, 13,198,448 high-quality SNPs were retained. Invariant sites were excluded at this stage to focus downstream analyses on polymorphic loci. The final high-quality SNP sets were merged using the MergeVcfs module from Picard Tools (v3.2.0).

### 3 Demographic inferences and recombination rate estimation

The demographic history of *H. sbordonii* and *H. semele* was reconstructed using multiple tools based on different modelling assumptions to increase the robustness and reliability of the results. SMC++, MSMC2 and Stairway Plot v2 were used to infer long-term changes in effective population size, while the LD-based method GONE was used to explore more recent demographic changes (within the last ∼200 generations), allowing us to test for recent declines that may have shaped the genomic landscape of *H. sbordonii*. For each population, we assumed a mutation rate of 2.9e⁻⁹ mutations per site per generation, based on estimates for *Heliconius melpomene* (Keightley et al. 2015).

#### 3.1 MSMC2

##### 3.1.1 Per species population size estimation

We inferred historical effective population size (Ne) trajectories and relative cross-coalescence rates (rCCR) using MSMC2 (v2.1.4) (Schiffels & Wang 2020). This analysis included the 10 *H. semele* mainland individuals, and 10 *H. sbordonii* Ponza individuals collected in 2019. Firstly, sex chromosomes were excluded from VCF files using vcftools v0.1.17. Next, to define genomic regions suitable for MSMC2, we created three masks: 1) a callable sites mask was generated for each individual bam file using GATK v3.8 CallableLoci, with a minimum base and mapping quality of 20, and coverage ranging between one-third and three times the individual’s mean depth (calculated using samtools v1.21 depth); 2) a mappability mask was generated from the reference genome with GenMap v1.3.0, retaining only positions with mappability ≥0.99; 3) a negative repeat mask was generated from the reference genome using RepeatMasker v4.1.7-p1, to exclude repetitive DNA and low-complexity regions. All output BED files were sorted with bedtools v2.32 sort. VCF files were phased with Beagle (default parameters with imputation enabled). As Beagle modifies VCF headers, we restored formatting using bcftools view and bcftools reheader. Files were compressed and indexed with bcftools. VCFs and mask files were split per sample and per chromosome, then formatted for MSMC2 using generate_multihetsep.py from MSMC-tools, applying all three masks. MSMC2 was run for each individual separately, and for all samples combined per species, with 20 replicates (-i 20) and the default time segment pattern (-p 1*2+25*1+1*2+1*3). Results were scaled assuming a mutation rate of 2.9e-9 per site per generation and a generation time of 1 year.

##### 3.1.2 Population divergence estimation (rCCR)

To assess population divergence, we performed a cross-population coalescence analysis. This requires all haplotypes to be present per chromosome, so the VCF files were merged across individuals using bcftools merge. A full analysis with 40 haplotypes (20 diploid individuals) was computationally unfeasible, so we used multihetsep_bootstrap.py from MSMC-tools to generate 20 bootstrap replicates, by sampling 5 Mb chunks with replacement into synthetic 4 Gb genomes. To further reduce memory usage, four haplotypes per species (eight total) were additionally subsampled randomly with replacement. MSMC2 was then run per species (e.g., haplotypes 1–4 for *H. sbordonii*, 21–24 for *H. semele*), and across all 16 cross-population haplotype pairs, by specifying haplotypes using the -I flag. To improve inflated Ne estimates at recent time points, we used a modified time segment pattern (-p 1*2+5*1+1*2). Outputs were merged using combineCrossCoal.py from MSMC-tools. Demographic trajectories and rCCR curves were visualized in Python using matplotlib, with cross-population rCCR summaries plotted as the median trajectory with shaded minimum and maximum ranges across replicates.

#### 3.2 SMC++ and recombination rate estimation with Pyrho

We inferred historical changes in effective population size using SMC++, which enables demographic inference from unphased genome data and can incorporate multiple individuals per population (Terhorst et al. 2017). Prior to analysis, the joint VCF file was filtered to exclude missing data, SNPs located within sex chromosomes, annotated gene regions, transposable elements (TEs), and low-mappability regions identified using GenMap (Pockrandt et al. 2020). Only autosomal scaffolds were retained. For each species, the five individuals with the highest sequencing coverage were selected to maximize genotype accuracy and ensure robust inference. These individuals were specified as *distinguished* samples in SMC++ and used to estimate the conditional site frequency spectrum (CSFS), with a thinning interval of 2000. In addition to reconstructing effective population size trajectories over time, we used SMC++ *split* to estimate the divergence time between *H. sbordonii* and *H. semele*. This analysis provided a timeframe for their evolutionary separation in the context of their demographic histories. Using the *Ne* trajectory after the split inferred for *H. sbordonii*, we estimated chromosome-specific recombination rates with Pyrho (Spence & Song 2019). To identify the best parameter combination for Pyrho’s optimization step, we first ran a hyperparameterization across multiple values of blockpenalty and windowsize. The final parameters (blockpenalty = 10, windowsize = 35) were selected based on composite likelihood and stability of inferred recombination-rate profiles across parameter combinations.

#### 3.3 Stairway Plot v2

We inferred past changes in effective population size (Ne) for *H. sbordonii* (recent samples only) and *H. semele* using Stairway Plot v2 (Liu & Fu 2020). To generate the required input site frequency spectra (SFS), we first processed the filtered VCF files of each species to exclude genomic regions that could bias demographic inference, such as regions under selection or miscalled regions. Specifically, we removed all sites located within sex chromosomes, annotated genes, transposable elements (TEs), and regions of low mappability identified using GenMap (Pockrandt et al 2020). Sites with missing genotype data across individuals were also excluded. We then computed the folded SFS for each population using easySFS (Gutenkunst et al. 2009). The resulting folded SFS files were used as input for Stairway Plot v2. A generation time of one year was used for both species, based on field observations and life cycle data. To estimate the total number of observed sites (required for Stairway Plot v2), we counted all callable sites, i.e., both polymorphic and monomorphic positions, after filtering out missing data and excluding genic, TE, and low-mappability regions. This ensured that both the SFS and site count were based on the same set of high-quality, neutrally evolving genomic regions. Stairway Plot v2 was then run independently for each population using recommended settings to assess confidence intervals around the estimated Ne trajectories.

#### 3.4 GONE

Following recombination rate estimation with Pyrho, we used the average chromosome recombination rate obtained for *H. sbordonii* as input for GONE, a method designed to infer recent demographic changes (<200 generations) from patterns of linkage disequilibrium (LD) (Santiago et al. 2020). GONE was applied to unphased genotypes from modern *H. sbordonii* individuals and *H. semele* samples using only high-quality SNPs located on autosomal scaffolds and excluding variants in gene regions, transposable elements, and low-mappability regions. We ran 20 independent replicates for both sample groups using default parameters and set the hc threshold to 0.01. This multi-replicate approach ensured robustness and allowed us to capture consistent signals of recent demographic dynamics.

### 4 Genomic diversity and population structure

To contextualize the levels of genomic diversity observed in our focal populations, we first estimated genome-wide heterozygosity in *H. sbordonii* and *H. semele* and compared these values with those of other Nymphalid butterfly species. To this end, we retrieved publicly available Illumina paired-end sequencing datasets (one per Nymphalid species) and processed them using a standardized workflow. Reads were quality-trimmed with fastp (Chen et al. 2018), and quality metrics were assessed using *FastQC* (Andrews et al. 2017). We then computed k-mer profiles using *Jellyfish* and estimated genome-wide heterozygosity with *GenomeScope2* (Ranallo-Benavidez et al. 2020), using a k-mer length of 21, an average read length of 150 bp, and a maximum k-mer count of 1000 to filter out repetitive elements. This comparative framework allowed us to place the genomic diversity of *H. sbordonii* and *H. semele* in the broader context of the Nymphalidae family. We next quantified genomic diversity across the dataset. For each population, nucleotide diversity (π) and population-level heterozygosity were calculated in non-overlapping 10 kb windows. Nucleotide diversity was estimated using VCFtools, whereas heterozygosity was calculated with a custom Python script from the https://github.com/sebafava/Hipparchia-sbodonii-population-genomics.git workflow based on the scikit-allel package. Additional population genetic statistics, including Tajima’s D and pairwise F*_ST_*, were likewise calculated in 10 kb windows using VCFtools. All analyses were performed on filtered VCF files restricted to autosomal chromosomes. To further explore population structure, we performed a pairwise relatedness analysis using VCFtools on filtered autosomal SNPs to detect potential cases of recent kinship among individuals. This analysis revealed that two *H. sbordonii* collected in Ponza in 2019 (i.e. individual 5 and 10) were strongly related with each other. Principal Component Analysis (PCA) was conducted using PLINK (Chang et al. 2015), SNPs were pruned for linkage disequilibrium and filtered to retain only autosomal chromosomes.

### 5 Inbreeding Estimation

To assess inbreeding levels and identify long tracts of homozygosity in the genome, we performed ROH detection using *BCFtools v1.21* (Narasimhan et al. 2016). The analysis was conducted on filtered, high-quality VCF files separately for each population group: *H. sbordonii* from Ponza (2019), *H. sbordonii* historical samples from Ponza and Palmarola (1990s), and *H. semele* from mainland Italy. Due to the lack of a detailed linkage map, we used physical distance as a proxy for recombination rates, assuming 1.57 cM/Mb, based on estimates obtained with *Pyrho*. We considered a ROH as a segment >50 kb with ≥100 markers and minimum average Phred score ≥30. As an independent validation, ROHs were also called from the joint dataset of complete biallelic autosomal SNPs using PLINK v1.9 (Chang et al. 2015), requiring a minimum length of 50 kb and 50 SNPs, a maximum average density of 50 kb per SNP, a maximum gap of 500 kb between consecutive SNPs, no more than one heterozygous site per ROH and per 50-SNP sliding window, no missing genotypes per window, and a window threshold of 0.05. To explore the distribution of ROH across individuals, we categorized ROH into five length classes (<100 kb, 100–250 kb, 250–500 kb, 500 kb–1 Mb, and >1 Mb). Genomic inbreeding for each sample was estimated from ROH (F_ROH_), calculated as the ratio of the total length of the genome covered by ROH to the total length of the genome covered by SNPs following the method proposed by McQuillan et al. (2008).

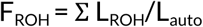

In which L_ROH_ is the total length of an individual’s ROH, and L_auto_ is the length of the autosomal genome covered by SNPs, which was 372.5 Mb. For each sample, five ROH estimates were calculated based on lengths from sequence data as the proportion of its genome: ROH<100 kb, 100–250 kb, 250–500 kb, 500 kb–1 Mb, and >1 Mb; F_ROH <100 kb_, F_ROH 100–250 kb_, F_ROH 250–500 kb_, F_ROH 500 kb–1 Mb_, and F_ROH >1Mb_.

### 6 Genetic load estimation

#### 6.1 Annotation-based approach to estimate the genetic load

To estimate the genetic load in *H. sbordonii* and *H. semele*, we employed an annotation-based strategy focused on coding sequence mutations. We started from a joint VCF dataset containing 13,198,448 SNPs and used SnpEff v5.1 to annotate variant effects. The chromosome-scale reference genome of *H. sbordonii* was used for annotation (Fava et al. 2024). Annotated genes were filtered using gFACs (Caballero & Wegrzyn 2019) to ensure compatibility with SnpEff and improve annotation accuracy. The filtering retained unique gene models with a minimum coding sequence length of 150 bp and removed incomplete transcripts at the 5′ and 3′ ends. The resulting annotation was used to build a custom SnpEff database. Variants were then categorized based on their predicted functional impact following SnpEff’s classification: “modifier,” “low,” “moderate,” and “high” effect (Cingolani 2012).

#### 6.2 Ancestral state definition

To avoid biases introduced by the reference genome and enable downstream analyses requiring polarized variants, we recoded genotypes based on the inferred ancestral state rather than relying on the reference allele. The ancestral state of annotated SNPs was defined using two outgroup species, *H. fagi* and *H. hermione*. We then retained SNPs that satisfied the following criteria: (1) both outgroup species and the two focal populations had non-missing genotype calls at the site, (2) the outgroups were fixed for the same allele, and (3) the two focal populations (*H. sbordonii* samples and *H. semele*) were not fixed for the same allele. These filters ensured that the sites represented informative variation relevant to the divergence and differentiation between the two species. Genotypes were then polarized so that outgroup individuals were consistently homozygous for the ancestral allele, ensuring uniform orientation across the dataset. When the ancestral allele matched the reference allele in the original VCF, genotypes were left unchanged; otherwise, alleles were systematically inverted across all samples. All these steps were automated through a custom Python script for filtering, allele comparison, and genotype re-coding (https://github.com/emitruc/genoloader.git). After retaining only the SNPs that followed the above criteria, 7,362,162 high-confidence polarized SNPs remained for downstream analyses. This approach minimized biases due to sequencing stochasticity and ensured more accurate comparisons of mutational load between populations.

#### 6.3 Proxies of relative genetic load and purging inference

Following the polarization of SNPs using outgroup-based ancestral state inference, we quantified the distribution of genotypes across different classes of mutational effects annotated with SnpEff. For each individual, we counted the number of homozygous ancestral, heterozygous, and homozygous derived genotypes separately for each impact category. These genotype counts were used as proxies to assess the genetic load in the two focal populations, *H. sbordonii* (recent samples) and *H. semele*. Specifically, we used the number of heterozygous sites in all the categories to estimate the masked load, representing deleterious mutations potentially hidden from selection due to heterozygosity. Conversely, the number of homozygous derived genotypes in these same categories was used to estimate the realized load, corresponding to mutations likely to affect fitness due to full expression of their effects. This approach provided a comparative framework for evaluating the accumulation and expression of deleterious mutations across the two species. Because populations with small effective sizes are more susceptible to the effects of genetic drift, we expect island populations to show an enrichment of fixed, deleterious alleles. To test this hypothesis, we calculated the R_XY_ and R′_XY_ statistics as described by Do et al. (2015). In addition to the interspecific comparison, we applied the same R_XY_ and R’_XY_ framework to contemporary Ponza samples from 2019 (X) and historical Ponza samples from 1990 (Y), using the same polarized SNP set to assess temporal changes in derived variation. Because this comparison involved ten contemporary but only three historical individuals, temporal estimates were interpreted cautiously due to the lower precision and statistical power associated with the historical sample. These statistics are designed to detect asymmetries in the number of derived alleles between two groups of genomes (X and Y) by quantifying the number of alleles present in one group but absent in the other. The statistics are formally defined as follows:

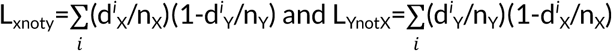

In this context, *d^i^* /*n* indicates the frequencies of derived alleles at site *i* in population X, while *d^i^* /*n* refers to the same quantities in population Y. In other words, we are quantifying how many derived mutations are present in population X but not in population Y, and vice versa. Based on these counts, we define the statistic R_XY_ as the ratio between them, which reflects the asymmetry in private derived variation between the two populations.

The ratio is computed as:

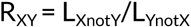

Because R_XY_ does not account for population substructure and demographic effects such as bottlenecks and inbreeding, we also computed R′_XY_, which normalizes the differences in frequency of derived alleles at functional sites using putatively neutral modifier variants as a reference (Do et al. 2015; Xue et al. 2015; Grossen et al. 2020).

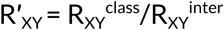

This correction helps disentangle the effects of natural selection from those of demographic history, allowing us to better detect signals of selection on functional variants. In this framework, R′_XY_ < 1 indicates a relative depletion of deleterious derived alleles in population X, consistent with effective purging of the genetic load in X compared to Y. Conversely, R′_XY_ > 1 suggests accumulation of deleterious mutations in population X. If natural selection has been equally efficient in both populations, R′_XY_ should approximate to 1. To explore the relative ratio of masked and realized loads, we also computed R_XY_ separately for homozygous and heterozygous genotypes, based on individual-level genotype calls. This approach allowed us to distinguish between the relative accumulation of derived alleles that are potentially expressed in homozygous states (realized load) and those likely masked in heterozygous states (masked load). For the homozygous-only analysis, we considered all sites where individuals were homozygous and calculated genotype frequencies in each population. Conversely, for the heterozygous-only analysis, we included only heterozygous genotypes.

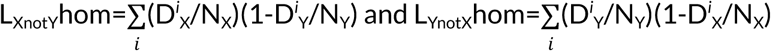

Where *D^i^_x_/N_x_* indicates the genotype frequencies of derived alleles in the homozygous state at site *i* in population X, while *D^i^_y_*/*N_y_* refers to the same quantities in population Y

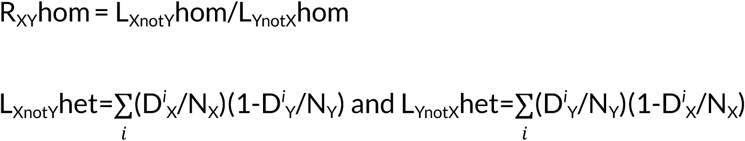

Where *D^i^_x_/N_x_* indicates the heterozygous genotype frequencies at site *i* in population X, while *D^i^_y_*/*N_y_* refers to the same quantities in population Y

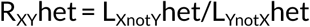

To obtain robust estimates and associated confidence intervals, we applied a weighted 100-block jackknife approach (Xue et al. 2015, Mathur et al. 2023; Grossen et al. 2020), following recent recommendations for the analysis of R_XY_ statistics (Stuart 2026). Variants were ordered by genomic position and divided into 100 contiguous blocks, after excluding SNPs located on the sex-linked scaffolds. R_XY_ and R′_XY_ were recalculated iteratively by leaving out one block at a time, and standard errors and confidence intervals were estimated using weighted block-jackknife equations. This procedure reduces linkage-related biases while avoiding the use of jackknife pseudo-values as independent observations, which can inflate the apparent precision of R_XY_ estimates.

### 7 Detection of candidate regions under selection and assessment of their contribution to R′_XY_ patterns

To identify putative regions under selection in *Hipparchia sbordonii* we combined three selection scans, SweepFinder2 (DeGiorgio et al. 2016), iHS and XP-EHH using the R package rehh (Gautier & Vitalis 2012), with window-based estimates of nucleotide diversity (π), Tajima’s D and F*_ST_*. SweepFinder2 candidates were defined as the top 1% of the genome-wide CLR distribution. For iHS, SNPs with -log_10_(p-value) > 3 were retained. For XP-EHH, only positive values were considered, indicating longer haplotypes in *H. sbordonii* relative to *H. semele*, and candidates were retained when -log_10_(p-value) > 3. Summary statistics were filtered by retaining windows in the upper 5% of F*_ST_*, the upper 5% of -log_10_(π), corresponding to the lowest π values, and both the lower and upper 5% tails of Tajima’s D.

Candidate SNPs from SweepFinder2, iHS and XP-EHH were converted into genomic intervals by adding 5 kb upstream and downstream of each selected position, and were then integrated with the 10 kb windows identified from π, Tajima’s D and F*_ST_*. All selected intervals from the six metrics (CLR, iHS, positive XP-EHH, Tajima’s D, low π and F*_ST_*) were combined into a single interval dataset. Overlapping intervals on the same chromosome were merged into broader candidate regions, and the presence or absence of each metric was recorded for every merged region. Regions were considered under selection when they were supported by at least five out of the six metrics.

To assess whether differences in R′_XY_ between *H. sbordonii* and *H. semele* may reflect signatures of positive selection, we implemented a block bootstrap approach comparing genomic regions inside and outside those identified as under selection as described above. All SNPs were first classified according to their genomic context (inside or outside regions under selection). Within each genomic context, SNPs were further partitioned by SnpEff functional class. To ensure comparability between genomic contexts, we randomly sampled three times as many sites from non-selected regions as were present within the regions under selection for each functional category. This approach provided a broader representation of the genomic background while maintaining comparability in the relative composition of functional effect classes. The resampling procedure was repeated 1,000 times, generating independent balanced datasets. For each genomic context at each iteration, SNPs were divided into 100 genomic blocks based on physical coordinates, and a leave-one-block-out procedure was applied to estimate R′_XY_ for each effect class.

To formally test for differences in R′_XY_ within and outside regions under selection, at each iteration we fitted a linear model with R′_XY_ as response variable and genomic context (inside vs outside regions under selection) as predictor, separately for low and moderate-effect categories. The analysis was restricted to low and moderate functional effect categories because these were the functional classes showing a detectable R′_XY_ excess and as variants in these classes are more likely to have medium functional impacts and therefore represent a plausible substrate for positive selection. The iterative approach allowed us to quantify the direction and magnitude of the association between genomic context and R′_XY_ while accounting for the uncertainty arising from stochastic variation in site sampling and block assignment. The distribution of regression coefficients across iterations was used to evaluate the robustness of any observed enrichment in R′_XY_ within regions under selection relative to the genomic background.

Genes overlapping candidate regions under selection were functionally annotated using BlastKOALA (Kanehisa et al. 2016) and KOBAS (Bu et al. 2021), with *Drosophila melanogaster* as the reference species. KOBAS was used to test functional over-representation against Gene Ontology, KEGG PATHWAY, and Reactome annotations using hypergeometric/Fisher’s exact tests, followed by Benjamini–Hochberg correction for multiple testing. Terms with an FDR-adjusted *P* < 0.05 were considered significant.

#### Use of artificial intelligence tools

AI-based language tools were used exclusively to improve grammar, clarity and readability. They were not used to generate data, perform analyses or determine scientific interpretations. All content and final wording were reviewed and approved by the authors.

## Data availability

The trimmed whole-genome resequencing reads generated in this study have been deposited in the NCBI Sequence Read Archive under BioProject accession PRJNA1444072. The RNA-seq reads used for the exploratory gene-expression analyses are publicly available from NCBI under BioProject accessions PRJNA1089943 for *H. sbordonii* and PRJEB50723 for *H. semele*.

## Code availability

All scripts used to perform the analyses and generate the results are available at https://github.com/sebafava/Hipparchia-sbodonii-population-genomics.git

## Acknowledgements

This paper and related research have been conducted during and with the support of the Italian inter-university PhD Programme in Sustainable Development and Climate Change (PhD-SDC; www.phd-sdc.it). This study was conducted within the framework of the ENDEMIXIT project, funded by the MUR (Italian Ministry for Research) PRIN 2017 grant 201794ZXTL and coordinated by the Department of Life Sciences and Biotechnology of the University of Ferrara (PI: Giorgio Bertorelle). Computational work was performed by the HPC cluster of the Department of Life and Environmental Sciences (DiSVA-HPC “ANNA.T”), Marche Polytechnic University

## Author Contributions

S.F. performed the main genomic analyses, contributed to the interpretation of the results, and led the writing of the manuscript. M.Ga. contributed to the design, calculation, block-bootstrap analysis, and interpretation of the purging analyses. D.K. performed the demographic reconstruction using MSMC2. P.G. contributed to supervision and interpretation of the results. D.C. proposed the initial study idea, provided the samples, and contributed to the interpretation of the results. A.I. performed DNA extraction, genomic library preparation, sequencing, and preliminary analysis of the raw sequencing data, and contributed to supervision and interpretation of the results. C.C. supervised DNA sequencing. R.B. contributed to bioinformatic analyses and supervision. M.Ge. contributed to supervision and interpretation of the results. G.B. contributed to the study design and supervised the analyses. E.T. performed DNA extraction for a subset of samples, designed the study, supervised the analytical design and interpretation of the results, and co-wrote the manuscript. All authors contributed to manuscript revision and approved the final version.

## References

Andrews, S. (2017, August). FastQC: a quality control tool for high throughput sequence data. 2010.

Beaurepaire, A. L., Webster, M. T., & Neumann, P. (2024). Population genetics for insect conservation and control. Conservation Science and Practice, 6(3), e13095.

Bertorelle, G., Raffini, F., Bosse, M., Bortoluzzi, C., Iannucci, A., Trucchi, E., … & Van Oosterhout, C. (2022). Genetic load: genomic estimates and applications in non-model animals. Nature Reviews Genetics, 23(8), 492–503.

Bonelli, S., Casacci, L. P., Barbero, F., Cerrato, C., Dapporto, L., Sbordoni, V., … & Balletto, E. (2018). The first red list of Italian butterflies. Insect Conservation and Diversity, 11(5), 506–521.

Bosse, M., & van Loon, S. (2022). Challenges in quantifying genome erosion for conservation. Frontiers in Genetics, 13, 960958.

Broquet, T., Angelone, S., Jaquiéry, J., Joly, P., Léna, J.-P., Lengagne, T., Plénet, S., Luquet, E., & Perrin, N. (2010). Genetic bottlenecks driven by population disconnection. Conservation Biology, 24(6), 1596–1605.

Bu, D., Luo, H., Huo, P., Wang, Z., Zhang, S., He, Z., … & Kong, L. (2021). KOBAS-i: intelligent prioritization and exploratory visualization of biological functions for gene enrichment analysis. Nucleic acids research, 49(W1), W317–W325.

Caballero, M., & Wegrzyn, J. (2019). gFACs: gene filtering, analysis, and conversion to unify genome annotations across alignment and gene prediction frameworks. Genomics, Proteomics & Bioinformatics, 17(3), 305–310.

Cardoso, P., Erwin, T. L., Borges, P. A., & New, T. R. (2011). The seven impediments in invertebrate conservation and how to overcome them. Biological conservation, 144(11), 2647–2655.

Cardoso, P., & Leather, S. R. (2019). Predicting a global insect apocalypse. Insect Conservation and Diversity, 12(4), 263–267.

Ceballos, F. C., Joshi, P. K., Clark, D. W., Ramsay, M., & Wilson, J. F. (2018). Runs of homozygosity: windows into population history and trait architecture. Nature Reviews Genetics, 19(4), 220–234.

Cesaroni, D., Lucarelli, M., Allori, P., Russo, F., & Sbordoni, V. (1994). Patterns of evolution and multidimensional systematics in graylings (Lepidoptera: Hipparchia). Biological Journal of the Linnean Society, 52, 101–119.

Chang, C. C., Chow, C. C., Tellier, L. C., Vattikuti, S., Purcell, S. M., & Lee, J. J. (2015). Second-generation PLINK: rising to the challenge of larger and richer datasets. Gigascience, 4(1), s13742–015.

Chávez, V. M., Marqués, G., Delbecque, J. P., Kobayashi, K., Hollingsworth, M., Burr, J., … & O’Connor, M. B. (2000). The Drosophila disembodied gene controls late embryonic morphogenesis and codes for a cytochrome P450 enzyme that regulates embryonic ecdysone levels. Development, 127(19), 4115–4126.

Chen, S., Zhou, Y., Chen, Y., & Gu, J. (2018). fastp: an ultra-fast all-in-one FASTQ preprocessor. Bioinformatics, 34(17), i884–i890.

Chowdhury, S., Dubey, V. K., Choudhury, S., Das, A., Jeengar, D., Sujatha, B., … & Kumar, V. (2023). Insects as bioindicator: A hidden gem for environmental monitoring. Frontiers in Environmental Science, 11, 1146052.

Chun, S., & Fay, J. C. (2011). Evidence for hitchhiking of deleterious mutations within the human genome. PLoS genetics, 7(8), e1002240.

Cingolani, P. (2012). Variant annotation and functional prediction: SnpEff. In Variant Calling: Methods and Protocols (pp. 289–314). New York, NY: Springer US.

Comay, O., Ben Yehuda, O., Schwartz-Tzachor, R., Benyamini, D., Pe’Er, I., Ktalav, I., & Pe’Er, G. (2021). Environmental controls on butterfly occurrence and species richness in Israel: The importance of temperature over rainfall. Ecology and Evolution, 11(17), 12035–12050.

Danecek, P., Auton, A., Abecasis, G., Albers, C. A., Banks, E., DePristo, M. A., … & 1000 Genomes Project Analysis Group. (2011). The variant call format and VCFtools. Bioinformatics, 27(15), 2156–2158.

Danecek, P., Bonfield, J. K., Liddle, J., Marshall, J., Ohan, V., Pollard, M. O., … & Li, H. (2021). Twelve years of SAMtools and BCFtools. Gigascience, 10(2), giab008.

De-Dios, T., Fontsere, C., Renom, P., Stiller, J., Llovera, L., Uliano-Silva, M., … & Lalueza-Fox, C. (2024). Whole genomes from the extinct Xerces Blue butterfly can help identify declining insect species. Elife, 12, RP87928.

DeGiorgio, M., Huber, C. D., Hubisz, M. J., Hellmann, I., & Nielsen, R. (2016). SweepFinder2: increased sensitivity, robustness and flexibility. Bioinformatics, 32(12), 1895–1897.

Dermauw, W., & Van Leeuwen, T. (2014). The ABC gene family in arthropods: comparative genomics and role in insecticide transport and resistance. Insect biochemistry and molecular biology, 45, 89–110.

Didham, R. K., Barbero, F., Collins, C. M., Forister, M. L., Hassall, C., Leather, S. R., … & Stewart, A. J. (2020). Spotlight on insects: trends, threats and conservation challenges. Insect Conservation and Diversity, 13(2), 99–102.

Díez-del-Molino, D., Sánchez-Barreiro, F., Barnes, I., Gilbert, M. T. P., & Dalén, L. (2018). Quantifying temporal genomic erosion in endangered species. Trends in Ecology & Evolution, 33(3), 176–185.

Do, R., Balick, D., Li, H., Adzhubei, I., Sunyaev, S., & Reich, D. (2015). No evidence that selection has been less effective at removing deleterious mutations in Europeans than in Africans. Nature Genetics, 47(2), 126–131.

Dunn, R. R. (2005). Modern insect extinctions, the neglected majority. Conservation Biology, 19(4), 1030–1036.

Dussex, N., Morales, H. E., Grossen, C., Dalén, L., & van Oosterhout, C. (2023). Purging and accumulation of genetic load in conservation. Trends in Ecology & Evolution, 38(10), 961–969.

Edwards, C. B., Zipkin, E. F., Henry, E. H., Haddad, N. M., Forister, M. L., Burls, K. J., … & Schultz, C. B. (2025). Rapid butterfly declines across the United States during the 21st century. Science, 387(6738), 1090–1094.

Fava, S., Sollitto, M., Racaku, M., Iannucci, A., Benazzo, A., Ancona, L., … & Trucchi, E. (2024). Chromosome-Level Reference Genome of the Ponza Grayling (*Hipparchia sbordonii*), an Italian Endemic and Endangered Butterfly. Genome Biology and Evolution, 16(7), evae136.

Forister, M. L., Pelton, E. M., & Black, S. H. (2019). Declines in insect abundance and diversity: We know enough to act now. Conservation Science and Practice, 1(8), e80.

Gautier, M., & Vitalis, R. (2012). rehh: an R package to detect footprints of selection in genome-wide SNP data from haplotype structure. Bioinformatics, 28(8), 1176–1177.

Gompert, Z., Springer, A., Brady, M., Chaturvedi, S., & Lucas, L. K. (2021). Genomic time-series data show that gene flow maintains high genetic diversity despite substantial genetic drift in a butterfly species. Molecular Ecology, 30(20), 4991–5008.

Grossen, C., Guillaume, F., Keller, L. F., & Croll, D. (2020). Purging of highly deleterious mutations through severe bottlenecks in Alpine ibex. Nature Communications, 11(1), 1001.

Gutenkunst, R. N., Hernandez, R. D., Williamson, S. H., & Bustamante, C. D. (2009). Inferring the joint demographic history of multiple populations from multidimensional SNP frequency data. PLoS Genetics, 5(10), e1000695.

Hallmann, C. A., Sorg, M., Jongejans, E., Siepel, H., Hofland, N., Schwan, H., … & De Kroon, H. (2017). More than 75 percent decline over 27 years in total flying insect biomass in protected areas. PLoS One, 12(10), e0185809.

Hill, P., Dickman, C. R., Dinnage, R., Duncan, R. P., Edwards, S. V., Greenville, A., Sarre, S. D., Stringer, E. J., Wardle, G. M., & Gruber, B. (2023). Episodic population fragmentation and gene flow reveal a trade-off between heterozygosity and allelic richness. Molecular Ecology, 32(24), 6766–6776.

Jangjoo, M., Matter, S. F., Roland, J., & Keyghobadi, N. (2016). Connectivity rescues genetic diversity after a demographic bottleneck in a butterfly population network. Proceedings of the National Academy of Sciences, 113(39), 10914–10919.

Jangjoo, M., Matter, S. F., Roland, J., & Keyghobadi, N. (2020). Demographic fluctuations lead to rapid and cyclic shifts in genetic structure among populations of an alpine butterfly, *Parnassius smintheus*. Journal of Evolutionary Biology, 33(5), 668–681.

Kanehisa, M., Sato, Y., & Morishima, K. (2016). BlastKOALA and GhostKOALA: KEGG tools for functional characterization of genome and metagenome sequences. Journal of molecular biology, 428(4), 726–731.

Kardos, M., Luikart, G., & Allendorf, F. W. (2015). Measuring individual inbreeding in the age of genomics: marker-based measures are better than pedigrees. Heredity, 115(1), 63–72.

Keightley, P. D., Pinharanda, A., Ness, R. W., Simpson, F., Dasmahapatra, K. K., Mallet, J., … & Jiggins, C. D. (2015). Estimation of the spontaneous mutation rate in *Heliconius melpomene*. Molecular Biology and Evolution, 32(1), 239–243.

Khan, A., Patel, K., Shukla, H., Viswanathan, A., van der Valk, T., Borthakur, U., … & Ramakrishnan, U. (2021). Genomic evidence for inbreeding depression and purging of deleterious genetic variation in Indian tigers. Proceedings of the National Academy of Sciences, 118(49), e2023018118.

Leung, K., Beukeboom, L. W., & Zwaan, B. J. (2025). Inbreeding and outbreeding depression in wild and captive insect populations. Annual Review of Entomology, 70(1), 271–292.

Lister, B. C., & Garcia, A. (2018). Climate-driven declines in arthropod abundance restructure a rainforest food web. Proceedings of the National Academy of Sciences, 115(44), E10397–E10406.

Liu, X., & Fu, Y. X. (2020). Stairway Plot 2: demographic history inference with folded SNP frequency spectra. Genome Biology, 21(1), 280.

Macdonald, Z. G., Dupuis, J. R., Glasier, J. R., Sissons, R., Moehrenschlager, A., Shaffer, H. B., & Sperling, F. A. (2025). Whole-genome evaluation of genetic rescue: the case of a curiously isolated and endangered butterfly. Molecular Ecology, 34(4), e17657.

Marsden, C. D., Ortega-Del Vecchyo, D., O’Brien, D. P., Taylor, J. F., Ramirez, O., Vilà, C., … & Lohmueller, K. E. (2016). Bottlenecks and selective sweeps during domestication have increased deleterious genetic variation in dogs. Proceedings of the National Academy of Sciences, 113(1), 152–157.

Mathur, S., Tomeček, J. M., Tarango-Arámbula, L. A., Perez, R. M., & DeWoody, J. A. (2023). An evolutionary perspective on genetic load in small, isolated populations as informed by whole genome resequencing and forward-time simulations. Evolution, 77(3), 690–704.

Mattila, A. L. K., Duplouy, A., Kirjokangas, M., Lehtonen, R., Rastas, P., & Hanski, I. (2012). High genetic load in an old isolated butterfly population. Proceedings of the National Academy of Sciences, 109(37), E2496–E2505.

McKenna, A., Hanna, M., Banks, E., Sivachenko, A., Cibulskis, K., Kernytsky, A., … & DePristo, M. A. (2010). The Genome Analysis Toolkit: a MapReduce framework for analyzing next-generation DNA sequencing data. Genome Research, 20(9), 1297–1303.

McQuillan, R., Leutenegger, A. L., Abdel-Rahman, R., Franklin, C. S., Pericic, M., Barac-Lauc, L., … & Wilson, J. F. (2008). Runs of homozygosity in European populations. The American Journal of Human Genetics, 83(3), 359–372.

Nabholz, B., Reboud, E. L., Lafon, B. J., Cotton, A. M., Partridge, M. G., Collins, N. M., & Condamine, F. L. (2026). Endemic but not eroded: Genomic distinctiveness and conservation genomics of the British swallowtail butterfly (Papilio machaon britannicus). Insect Conservation and Diversity.

Narasimhan, V., Danecek, P., Scally, A., Xue, Y., Tyler-Smith, C., & Durbin, R. (2016). BCFtools/RoH: a hidden Markov model approach for detecting autozygosity from next-generation sequencing data. Bioinformatics, 32(11), 1749–1751.

Niwa, R., & Niwa, Y. S. (2014). Enzymes for ecdysteroid biosynthesis: their biological functions in insects and beyond. Bioscience, biotechnology, and biochemistry, 78(8), 1283–1292.

Nolen, Z. J., Rundlöf, M., & Runemark, A. (2024). Species-specific erosion of genetic diversity in grassland butterflies depends on landscape land cover. Biological Conservation, 296, 110694.

Pockrandt, C., Alzamel, M., Iliopoulos, C. S., & Reinert, K. (2020). GenMap: ultra-fast computation of genome mappability. Bioinformatics, 36(12), 3687–3692.

Purfield, D. C., Berry, D. P., McParland, S., & Bradley, D. G. (2012). Runs of homozygosity and population history in cattle. BMC Genetics, 13, 1–11.

Ranallo-Benavidez, T. R., Jaron, K. S., & Schatz, M. C. (2020). GenomeScope 2.0 and Smudgeplot for reference-free profiling of polyploid genomes. Nature Communications, 11(1), 1432.

Ringbauer, H., Novembre, J., & Steinrücken, M. (2021). Parental relatedness through time revealed by runs of homozygosity in ancient DNA. Nature Communications, 12(1), 5425.

Robinson, J., Kyriazis, C. C., Yuan, S. C., & Lohmueller, K. E. (2023). Deleterious variation in natural populations and implications for conservation genetics. Annual Review of Animal Biosciences, 11(1), 93–114.

Salvador, R. B., Tomotani, B. M., O’Donnell, K. L., Cavallari, D. C., Tomotani, J. V., Salmon, R. A., & Kasper, J. (2021). Invertebrates in science communication: confronting scientists’ practices and the public’s expectations. Frontiers in Environmental Science, 9, 606416.

Sánchez-Bayo, F., & Wyckhuys, K. A. (2019). Worldwide decline of the entomofauna: A review of its drivers. Biological Conservation, 232, 8–27.

Santiago, E., Novo, I., Pardiñas, A. F., Saura, M., Wang, J., & Caballero, A. (2020). Recent demographic history inferred by high-resolution analysis of linkage disequilibrium. Molecular Biology and Evolution, 37(12), 3642–3653.

Schiffels, S., & Wang, K. (2020). MSMC and MSMC2: the multiple sequentially Markovian coalescent. In Statistical Population Genomics (pp. 147–165). Humana.

Spence, J. P., & Song, Y. S. (2019). Inference and analysis of population-specific fine-scale recombination maps across 26 diverse human populations. Science Advances, 5(10), eaaw9206.

Stuart, O. P. (2026). On the Use of the Jackknife to Compare R_XY_ Statistics in Conservation Genomics Research. Molecular Ecology Resources, 26(4), e70148.

Sucháčková Bartoňová, A., Linke, D., Klečková, I., de G. Ribeiro, P., & Matos-Maraví, P. (2023). Incorporating genomics into insect conservation: Butterflies as a model group. Insect Conservation and Diversity, 16(4), 427–440.

Terhorst, J., Kamm, J. A., & Song, Y. S. (2017). Robust and scalable inference of population history from hundreds of unphased whole genomes. Nature Genetics, 49(2), 303–309.

Trucchi, E., Massa, P., Giannelli, F., Latrille, T., Gargano, M., Nitta Fernandes, F. A., … & Le Bohec, C. (2025). High gene expression predicts extremely low segregation of deleterious mutations in large penguin populations. Molecular biology and evolution, 42(6), msaf146.

Vasimuddin, M., Misra, S., Li, H., & Aluru, S. (2019, May). Efficient architecture-aware acceleration of BWA-MEM for multicore systems. In 2019 IEEE International Parallel and Distributed Processing Symposium (IPDPS) (pp. 314–324). IEEE.

Wagner, D. L., Grames, E. M., Forister, M. L., Berenbaum, M. R., & Stopak, D. (2021). Insect decline in the Anthropocene: Death by a thousand cuts. Proceedings of the National Academy of Sciences, 118(2), e2023989118.

Wang, Y., Allen, S. L., Reddiex, A. J., & Chenoweth, S. F. (2024). The impacts of positive selection on genomic variation in Drosophila serrata: Insights from a deep learning approach. Molecular Ecology, 33(18), e17499.

Warren, M. S., Maes, D., van Swaay, C. A., Goffart, P., Van Dyck, H., Bourn, N. A., … & Ellis, S. (2021). The decline of butterflies in Europe: Problems, significance, and possible solutions. Proceedings of the National Academy of Sciences, 118(2), e2002551117.

Webster, M. T., Beaurepaire, A., Neumann, P., & Stolle, E. (2023). Population genomics for insect conservation. Annual Review of Animal Biosciences, 11(1), 115–140.

Whitla, R., Hens, K., Hogan, J., Martin, G., Breuker, C., Shreeve, T. G., & Arif, S. (2024). The last days of *Aporia crataegi* (L.) in Britain: evaluating genomic erosion in an extirpated butterfly. Molecular Ecology, 33(19), e17518.

Xue, Y., Prado-Martinez, J., Sudmant, P. H., Narasimhan, V., Ayub, Q., Szpak, M., … & Scally, A. (2015). Mountain gorilla genomes reveal the impact of long-term population decline and inbreeding. Science, 348(6231), 242–245.

